# Bridging forward-in-time and coalescent simulations using pyslim

**DOI:** 10.1101/2025.09.30.679676

**Authors:** Shyamalika Gopalan, Murillo F. Rodrigues, Peter L. Ralph, Benjamin C. Haller

## Abstract

Ancestral Recombination Graphs (ARGs) provide an expressive and compact way to represent genetic variation data generated by simulations embedded within its genealogical history, and it can dramatically speed up simulations. The fact that the ARG records genealogical as well as genomic information opens up the possibility for a number of new analysis and simulation techniques. Here, we aim to introduce the reader to this deep source of information as produced by the forwards simulator `SLiM`. `SLiM` records the ARG using the *tree sequence* format, which can be manipulated using the `tskit` and `pyslim` python packages. We first describe the information that `SLiM` records in the tree sequence, then provide several examples that use the tree sequence as a format to losslessly pass population states between simulators: *recapitation* of a forwards simulation with coalescent simulation; initiation of a forwards simulation using the results of a coalescent simulation; and parallelization of simulations, first across branches of a phylogenetic tree, and then across the populations of parasites infecting different hosts.

## Introduction

Simulations have become an invaluable tool in population genetics over the past six decades, allowing researchers to model increasingly complex evolutionary scenarios. A robust research community has grown around developing computational methods for conducting these simulations, as well as for representing and analyzing the data that they produce.

In this chapter, we present a practical introduction to `pyslim`, a Python package that facilitates *hybrid simulations* using `SLiM` and `msprime`. Hybrid simulations are frequently used to combine key features of the two main methods of individual-based population genetic simulation. These two methods differ primarily in the direction of the simulation process – either forward- or backward-in-time. The coalescent process models the ancestry of sampled genomes backward-in-time until they coalesce into their most common recent ancestor. This approach is extremely efficient because it only simulates the ancestors that directly contributed to the sampled genomes, thus avoiding the need to represent large swathes of the population pedigree. The downside of this approach is its strict assumptions (e.g., neutrality, random mating within subpopulations), which limit applicability. On the other hand, forward-in-time simulations apply demographic processes such as reproduction, migration and mutation beginning with a set of ancestral individuals representing the entire population. This affords forward-in-time simulations much more flexibility, but at a higher computational cost.

Hybrid simulations have recently emerged as a strategy for leveraging the benefits of both forward-in-time and coalescent methods, as well as parallel computing, to efficiently model highly complex evolutionary scenarios. Many forward-in-time simulations conducted using `SLiM`, for example, are later modified using the coalescent process to ensure the coalescence of the individuals in the first generation, a process known as *recapitation* [Kelleher et al., 2018], which will be discussed in more detail later in this chapter. Since recapitation provides the genetic diversity of the initial generation it can be particularly helpful in expensive simulations – for instance, in the spatial simulations of Battey et al. [2020] and Petr et al. [2023]. As described later in this chapter, Rodrigues et al. [2024] performed whole-chromosome simulations of 10 million years of the entire great apes history, which would not have been possible without hybrid simulation techniques.

Hybrid simulations would not be possible without Ancestral Recombination Graphs (ARGs), a general class of data structures reviewed in Wong et al. [2024]. Compared to genotypes, the ARG represents a much richer source of information about the processes that gave rise to a given set of genomes. There are different formats for working with ARGs [Palamara, 2016, DeHaas et al., 2024, Parida et al., 2011, Gunnarsson et al., 2024]; here, we use the tree sequence data structure [Kelleher et al., 2016, Ralph et al., 2020] that is used by `msprime` and `SLiM` via the C and python libraries provided by `tskit`.

Here, we will review how `SLiM`, a popular forward-in-time simulation software, uses the tree sequence before describing the main uses of `pyslim` in conducting hybrid simulations, specifically: (i) recapitation, the process of simulating the history of uncoalesced first-generation individuals, (ii) generation of initial diversity using the coalescent process for forward-in-time simulations with selection, (iii) parallelization of multi-population simulations. Finally, to illustrate the realism that can be achieved with a hybrid approach, we present a vignette of an organism evolving under a complex life history. For further details on these examples, refer to the `pyslim` documentation (https://tskit.dev/pyslim/docs/latest), from which some of the text presented here was adapted. The code used in this paper is available in the accompanying Jupyter notebooks at https://github.com/petrelharp/pyslim_paper/tree/main/code.

## 1 Tree sequences and SLiM

An “ancestral recombination graph” (ARG) represents paths of genealogical inheritance and mutation that have produced a given set of genomes, and the *tree sequence* format of `tskit` provides a general-purpose way to store ARGs. See Lewanski et al. [2024] and Brandt et al. [2022] for reviews of the applications of ARGs and Wong et al. [2024] for an overview of the terminology and history of ARGs as we use it here. For this reason, we mostly use the term “tree sequence” in this chapter even where “ARG’ would be equally accurate, as well as other terminology associated with `SLiM` and `tskit` [Kelleher et al., 2016, Ralph et al., 2020, Wong et al., 2024]. Other simulators also produce tree sequences, but we have not tested those with `pyslim`, and so do not discuss them here.

As `SLiM` proceeds with simulating genomes evolving forward-in-time, if “tree sequence recording” is enabled, it tracks how all the genomes are related to each other and returns the result as an ARG in tree sequence format. First, we describe *what* information is recorded, *how* it is recorded, and how to access it.

### Terminology

Each *node* of a tree sequence represents a single (haploid) genome; inheritance relationships between these are recorded as *edges*, and genetic variation is represented by *mutations* at associated *sites*. A particular tree sequence describes the entire genealogy of a set of “focal” nodes called *sample nodes*, or simply *samples*, over the simulated time period. Many `tskit` operations act on these sample nodes by default. The samples are those genomes for which we have complete genetic ancestry information; we likely only have partial information about the other ‘non-sample’ nodes in the tree sequence, which are included because they are required to describe that ancestry. (“Who is and isn’t in the tree sequence” below gives more detail about sample nodes.) By default, `SLiM` simulates diploid organisms, where each *individual* ‘s genome is represented by two nodes.

Since `SLiM` is a forwards simulator, it records time (in units called “ticks”) since the start of the simulation, with 0 corresponding to the earliest time point. We call this `SLiM` *time*. However, in many cases, a more natural point of reference might be in relation to the samples (which are after all probably “sampled” today), and so time in the tree sequence is measured in units of *time ago*. This is the same way that time is represented in coalescent or backward-in-time approaches, with a time of 0 corresponding to the end of the simulation rather than the beginning. This dual representation of time is important to keep in mind when thinking about hybrid simulations, as we will see later in the chapter.

### Who is and isn’t in the tree sequence

Suppose we have run a very small simulation with `SLiM`. The genetic relationships among each of the diploid individuals who were alive over the course of the simulation might look something like Figure 1A. Since individuals (circles) are diploid, each contains two genomes or nodes (shaded rectangles). The edges of the tree sequence record which specific parts of the genome are inherited, so the relationships recorded in the edges of a tree sequence are between the nodes and not the individuals directly.

**Figure 1.**
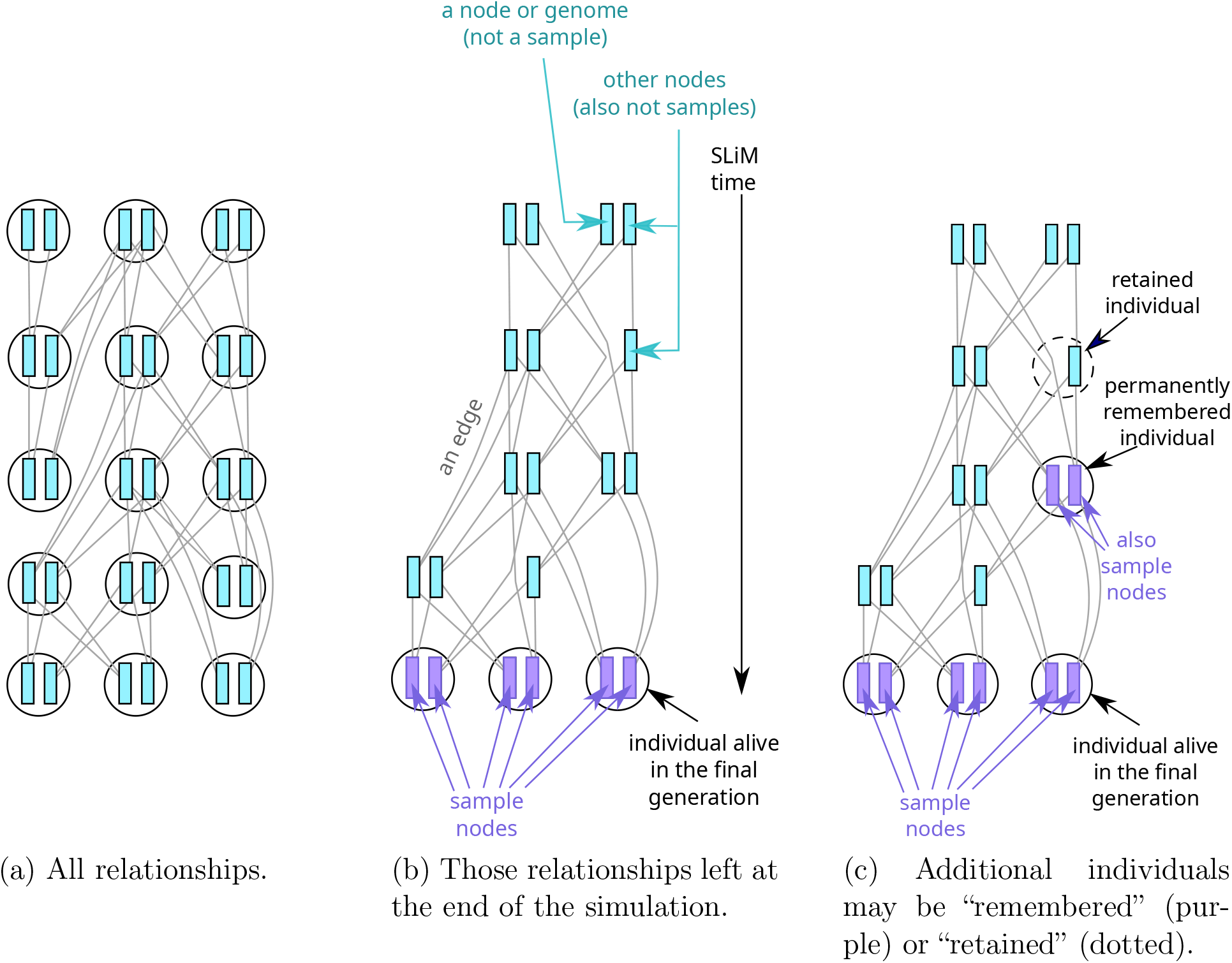
The resulting genetic relationships between individuals in a small simulation (three diploids over five generations).

By default, the sample nodes are simply the genomes of the individuals that are alive in the final generation of the simulation. This renders large portions of these relationships unnecessary for representing the samples’ history, as they correspond to lineages that died out at some earlier point in the simulation. To avoid having to store this large quantity of unnecessary information, `SLiM` periodically *simplifies* the tree sequence as the simulation goes along. By default, simplification will only keep the portions of the genetic genealogy that are required to represent the history of the current nodes [Kelleher et al., 2018]. Additionally, it defaults to removing any node that is not a genetic *most recent common ancestor* (MRCA) of at least two daughter nodes. This removes historical nodes that are only “on the line to” a sample, but do not represent a branching point (i.e., coalescent event) on the genealogical tree at some point on the genome. Below, we discuss how to retain additional historical information.

As a result of the simplification process, the output tree sequence will look something like Figure 1B. To be precise, edges correspond to inherited genomic *segments* with defined start and end positions, but we do not attempt to depict that complexity here.

Individuals who are alive at the end of the simulation automatically have their nodes marked as *samples. No* other individuals (depicted as circles) are present, although some of their nodes remain. In `SLiM`, certain information (including spatial location and age) is associated with individuals, not nodes, so by default we do not have access to this information for historical individuals. `tskit` also allows individuals to have a “parents” property that `SLiM` makes use of, so parentage information is also available even if no genetic material has been transmitted, but only if both parent and child individuals are represented in the tree sequence.

## Historical individuals

By default, only the nodes associated with individuals alive in the final generation are part of the set of samples. However, there are many cases where we might want to retain the complete ancestry, and other metadata, for historical individuals. For example, we might want to model the relationship between a modern population and one particular individual from the past. Or, as below, if we are conducting simulations in parallel that share a common ancestry, we may need to retain certain nodes that are critical for linking distinct tree sequences back together. In order to accomplish this, we can choose to “remember” key individuals during the course of the simulation, using the `SLiM` function `treeSeqRememberIndividuals()`.

### Permanently remembering individuals

By default, a call to `treeSeqRememberIndividuals()` will permanently remember one or more individuals, the simulated equivalent of ancient DNA dug out of permafrost. This means that any tree sequence subsequently output will always contain this individual, its nodes (now marked as samples), and its full ancestry. The result of remembering an individual in the introductory example is pictured in Figure 1C. This is also useful to, for instance, access allele frequencies and spatial locations of individuals at a particular time in the past. As an extreme case, all individuals at all time points can be remembered, thus retaining the complete population pedigree of everyone ever alive, but this quickly becomes computationally burdensome.

### Retaining individuals

Alternatively, you may want to only retain historical individuals as long as their nodes are still relevant to reconstructing the genetic ancestry of the sample nodes. The ancestral nodes that `SLiM` includes in the tree sequence do not by default have their associated individuals included as well, and so at the end of the simulation we do not by default have access to individual-level information for non-sample nodes such as spatial location or parental IDs. But, we can ask `SLiM` to also record historical individuals (and hence their associated information) as long as their nodes are retained through simplification. You can *retain* individuals in this way by using `treeSeqRememberIndividuals(…, permanent=F)`. This is less burdensome than permanently remembering them because individuals can still be removed by simplification once their nodes (which are not marked as samples) are no longer ancestral to the samples. Since a retained individual’s nodes are not samples, they are subject to the standard removal ‘rules’ of simplification. It is therefore possible to end up with an individual containing only one genome, as shown in the diagram. However, as soon as both nodes of a retained individual have been lost, the individual and any information associated with it is deleted as well.

As previously discussed, simplification will, by default, only keep nodes if they are a coalescent point (i.e., they are a MRCA or branch point) somewhere along the genome. This can be changed by initialising tree sequence recording in `SLiM` using `treeSeqInitialize(retainCoalescentOnly=F)`. `SLiM` will then preserve all retained individuals while they remain in the genealogy of present-day individuals, even if their nodes are not coalescent points in the tree. If you later decide to reduce the number of samples in the tree sequence, you can do so using the tskit function `simplify()` (which is what `SLiM` uses under the hood). In this case, individuals that are “retained” rather than “remembered” will be kept only if they are still MRCAs in the ancestry of the selected samples. This behavior corresponds to the `keep_unary_in_individuals` argument to `tskit.simplify`.

## 2 Recapitation: tying up loose ends

By default, a `SLiM` simulation begins with all individuals being identical, and genetic diversity builds up over time as new mutations occur. However, in many cases, starting with a clonal population is undesirable and can have long-lasting effects on patterns of diversity in the samples. One way to overcome this is to include a long “burn-in” period in the forward simulation to reach an equilibrium level of genetic diversity. However, this period generally needs to be quite long (on the order of 5–15 times the effective population size, in generations), which can be prohibitive. An efficient alternative is to “seed” the simulation with genetic diversity generated by a coalescent simulation, i.e., conduct a hybrid simulation. There are two main ways to do this, which we refer to as “recapitation” and “generating initial diversity”, respectively. Both ways should be thought of as generating initial diversity, but recapitation (counterintuitively) generates that diversity after the fact. So, the main difference between them is when this coalescent step is run relative to the forward-in-time portion of the simulation. We will discuss these approaches in the following two sections.

Recapitation is done with a tree sequence generated by `SLiM` simulation. This is suitable if the `SLiM` step does not require the ancestral genotypes for anything, such as determining individual fitness, so all we want to know is how the demographics of the `SLiM` simulation affects initially present neutral variation. Functionally, recapitation conducts a coalescent simulation starting with those portions of the first-generation ancestral nodes (`SLiM` time 0) that have not yet coalesced. This is discussed in more detail in Haller et al. [2019].

Figure 2 illustrates this process: imagine that, at some sites, one or more samples do not share a common ancestor within the `SLiM` simulated portion of history (shown in blue). Recapitation starts at the *top* of the genealogies and moves backwards in time to fill in a genealogical history for all the samples. The green chromosomes here are new ancestral nodes that have been added to the tree sequence. As previously mentioned, the effect of omitting this step would be a genetically homogeneous initial population; our simulation would have less genetic variation than it “should” have, since all the variation contributed by the green portion of the tree would be omitted.

**Figure 2.**
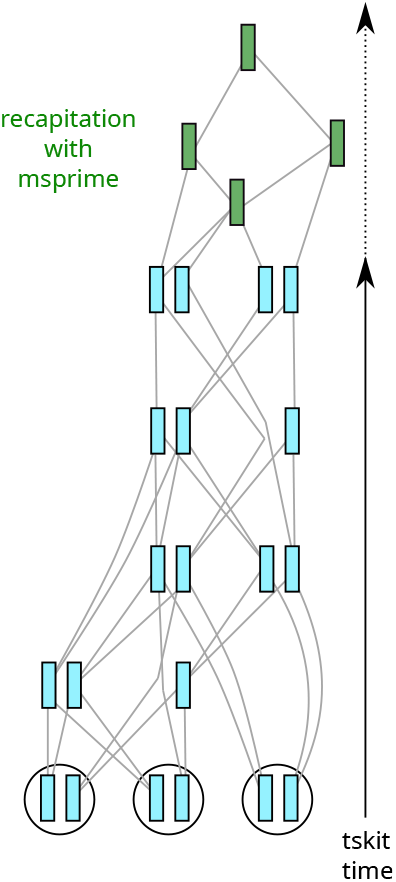
Recapitation adds the green nodes to the simulation of Figure 1 by coalescent simulation.

Recapitation within a single randomly-mating population of a size `N` can be achieved with a simple call to `pyslim.recapitate(ts, ancestral_size=N) Since recapitate` is a wrapper around `msprime.sim_ancestry`, we could recapitate our tree with any model that can be represented by `msprime` code [Baumdicker et al., 2022]. For instance, it is possible to recapitate a `SLiM`-generated tree sequence with a fluctuating population size or non-uniform genetic map by simply passing in the relevant arguments.

Setting up a demographic model in `msprime` that is consistent and compatible with a particular `SLiM` simulation requires understanding some of the finer details of each of these software packages, so here we will present a concrete example. Suppose our `SLiM` model has the following population structure:

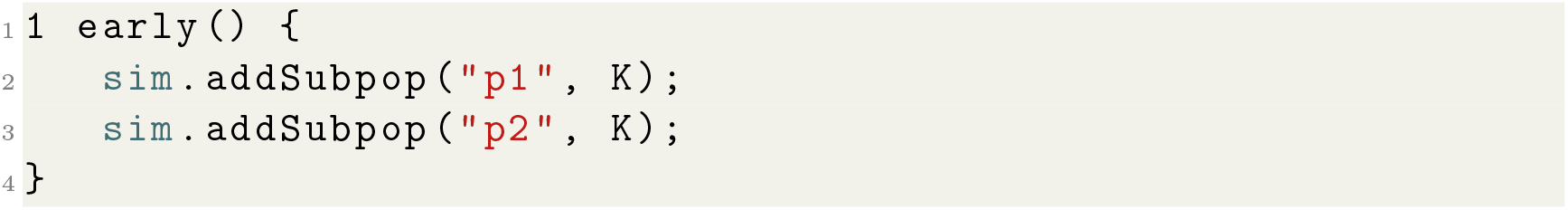

for some population size `K`. If the resulting tree sequence is stored in the file `recap_example.trees`, then examining the populations using the shell, we see (edited to remove some extraneous information):

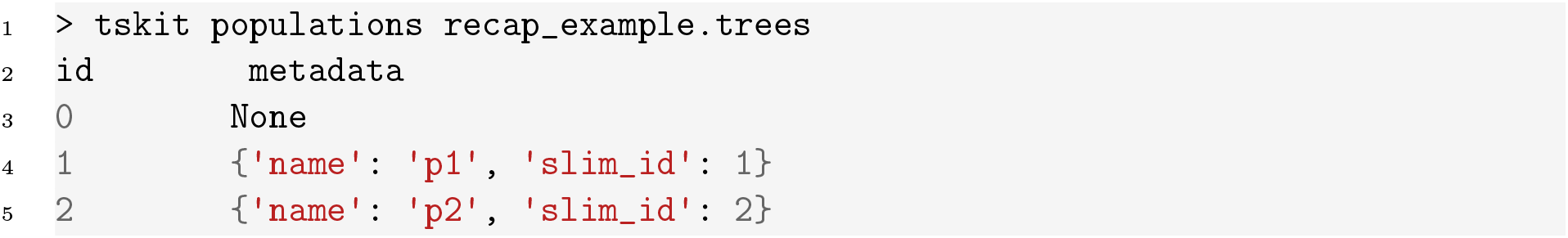

(We could see this in python by `print`ing the relevant population objects.) There are *three* populations, but the first one is unused (and has no metadata), because `SLiM` stores population `pX` as population `X`, and our `SLiM` script had subpopulations `p1` and `p2`. To be compatible with this tree sequence, the `msprime` demographic model will need to include three populations as well. However the first population will remain unused unless we specify it as an ancestral source for one of the other two populations during the coalescent phase of the simulation.

Regardless of the population numbering scheme used, `msprime` provides a built-in method of seamlessly generating a compatible demography given a particular starting tree sequence file. The following code uses the information in the tree sequence `ts` to create a basic compatible demography:

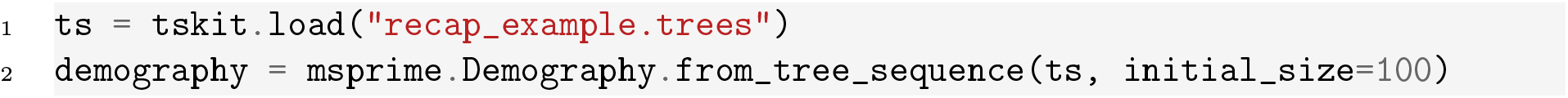

The resulting demography has 3 populations with an effective size of 100 individuals each. Note that the `initial_size` does not have to match what we specified in our `SLiM` simulation, because it specifies the size of the generation immediately preceding the `SLiM` phase. We could theoretically try to recapitate our tree sequence with this demography, but if we did so, `msprime` would run forever. This is because we have not allowed for any way for lineages from population `p1` to coalesce with lineages in population `p2`, or vice versa. Therefore, the coalescent process will never be able to conclude.

To remedy this, let us specify one-directional migration from `p1` to `p2` (so, lineages in `p2` move into `p1` as we move backwards in time) starting 100 generations before the start of the `SLiM` simulation. The current time (“tick”) in `SLiM` is stored in the tree sequence’s top-level metadata, as `ts.metadata[`‘`SLiM`’][‘`tick`’], which we can use to identify when the desired time point is (and note that this must be specified in tskit time, i.e., as “time ago”).

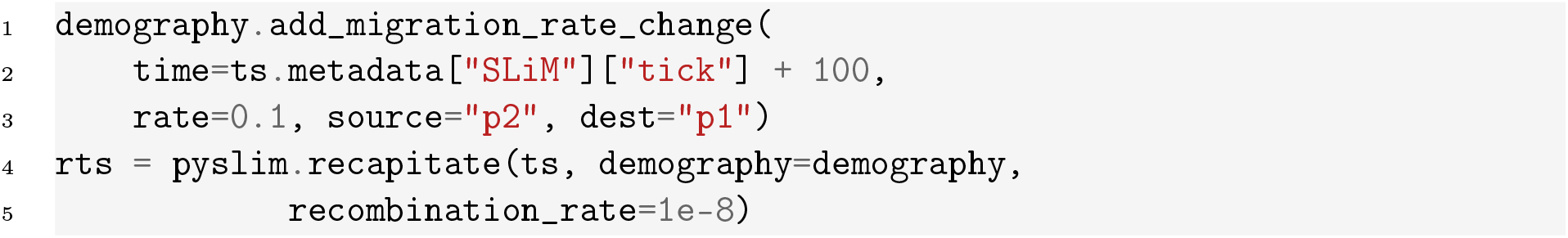

The `pyslim` function ‘recapitate’ passes the tree sequence `ts` to the `initial_state` argument of `msprime.sim_ancestry`, along with the demographic model that we specified. The root of each uncoalesced lineage in this tree sequence is a node that exists at a particular time and in a particular population; these two things situate the lineage within the demographic model, allowing `msprime` to continue following the lineages back through time.

At this point we need to raise an important caveat: in non-Wright-Fisher models in `SLiM`, time units are likely not in units of generations. This means that *different* mutation and recombination rates need to be passed to `msprime`, because these rates in `SLiM` are “per generation” while for `msprime` they are “per unit time”. For these and other considerations, see the `pyslim` documentation.

## 3 Generating initial diversity

If the `SLiM` simulation needs to actually use the genotypes of the initial generation, then we cannot recapitate. Instead, we can run the coalescent simulation first to generate a starting tree sequence, which is then loaded into `SLiM` where the main simulation process can proceed as normal. This approach is useful, for instance, when simulating selection on standing variation in a large population, where a purely forward-in-time burn-in period would be very costly in terms of time and computational resources. While running a neutral coalescent burn-in is not exactly the same as running a forward-in-time burn-in with selection, this can be an adequate approximation (but caution is advised).

For instance, imagine that we want to perform a lab experiment in which we take high-diversity organisms from the wild and subject them to selection for a few dozen generations. If we use the results of a coalescent simulation to represent our sample from the wild, then we are effectively assuming that the genetic diversity in the wild population is (approximately) neutral. This may not reflect reality: for instance, alleles with larger effect on a trait under stabilizing selection might be expected to be at lower frequency. For these reasons, we still suggest some burn-in in `SLiM`, although less is probably required, since you don’t have to wait for diversity to be generated by mutation. For our example, we will assume simply that the trait we are selecting for in the lab was not under selection in the wild. To implement this, we will: (1) run a coalescent simulation with `msprime`; (2) “annotate” the resulting tree sequence with `SLiM` metadata, allowing it to be read into `SLiM`; (3) add `SLiM` mutations with `msprime` and edit the mutation metadata to assign selection coefficients; and finally (4) run the `SLiM` portion of the simulation.

### Annotating a tree sequence with SLiM metadata

Suppose we run the following coalescent simulation in `msprime`:

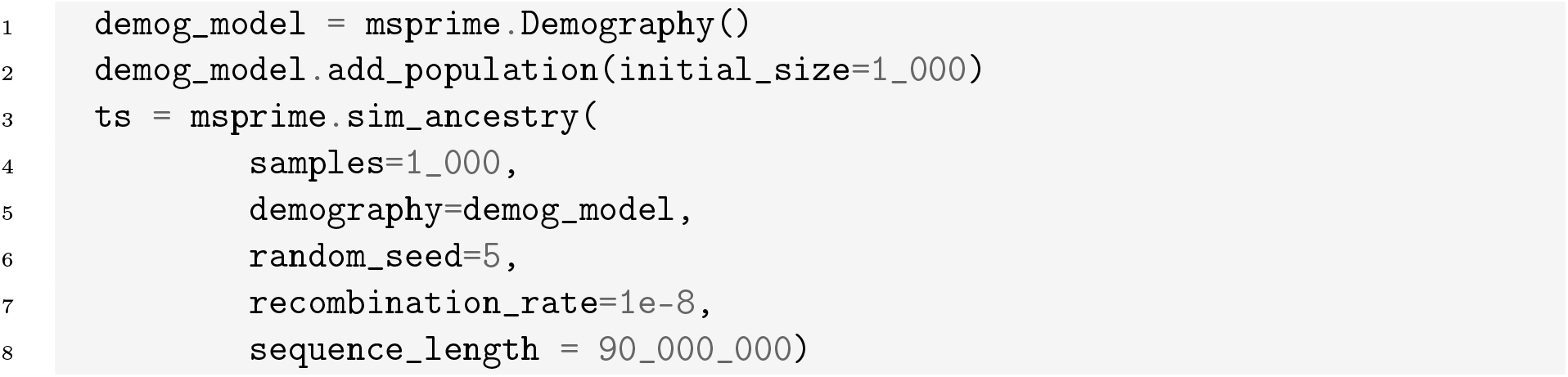

This tree sequence `ts` is a complete record of the genealogical information relating specific individuals and nodes, but is missing other information that `SLiM` requires. The other information needs to be added to metadata of various objects in the tree sequence, and can be easily added using the `annotate` function from `pyslim`. Although this function adds default values for most information, we must provide the model type (Wright-Fisher or non-Wright-Fisher) and some information about how coalescent- and `SLiM`-time should be aligned.

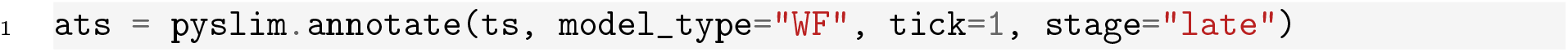

When this tree sequence is loaded into `SLiM`, the tick counter (i.e., `SLiM` time) will be set to 1, and if it is not loaded in during the `late()` stage a warning will be raised.

The function `annotate()` fills in default values for every element of the tree sequence that is required by `SLiM`, which includes all individuals, all nodes that exist within living individuals, and all populations referenced by nodes. We can see exactly what these defaults are for any given component of the tree sequence by using `default_slim_metadata`, for instance:

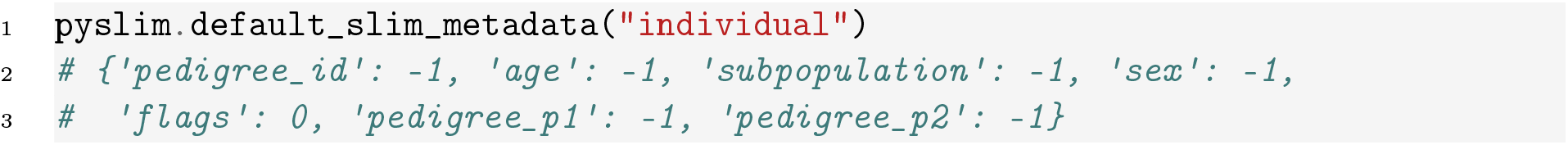

So, by default all individuals are hermaphroditic (individual `sex` is −1) and have `age` equal to −1 (as appropriate for a WF simulation). If we wanted, for instance, a sexual population we would then edit these values. Below, we show how to edit the metadata of mutations; other modifications such as modifying individuals can be done in a similar manner.

### Adding mutations with SLiM metadata

The simulation thus far had no mutations, so the next step is to generate the initial genetic diversity that natural selection will act on in the forward-in-time phase of our hybrid simulation. Let us assume an overall rate of 3 ×10^−8^ new mutations per base pair per generation. Further, let us assume that 99% of all new mutations are neutral, and the remaining 1% have a selection coefficient of 0.001. If we were doing this simulation entirely within a forward-in-time framework with a long burn-in, we could have achieved this outcome using the following `SLiM` code:

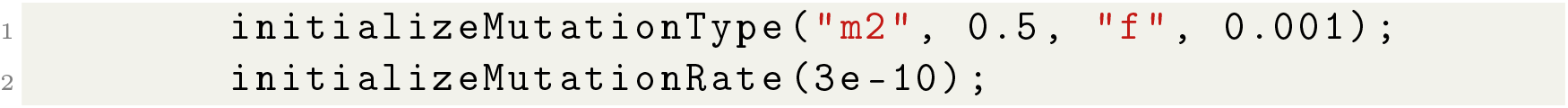

This code defines a mutation type `m2` that has a dominance coefficient of 0.5 and a fixed selection coefficient of 0.001 (the latter can be made more flexible, e.g., drawn from a distribution – see the `SLiM` manual). Notice, too, that we have set these non-neutral mutations to arise at a rate of 3 × 10^−10^ per base per generation, or 1% of our desired overall mutation rate of 3 × 10^−8^, so we would add the remaining neutral mutations after the fact (as we do below here as well). To approximate this result using `msprime`, we must first use the function `msprime.sim_mutations` with the `SLiMMutationModel` to generate some mutations on the existing tree sequence:

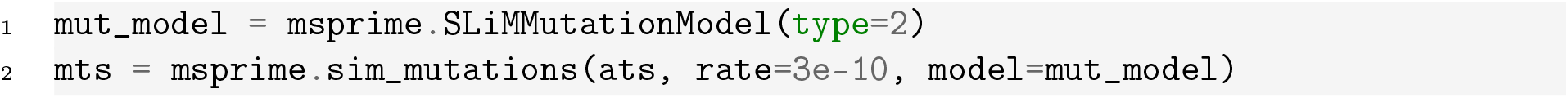

The `type=2` argument denotes that the mutations will be of type `m2` in `SLiM`, which will also need to be explicitly initialized in the forward-in-time phase of the simulation (as in the `SLiM` code above). By default, these mutations are assigned a selection coefficient of 0, which is accurate since they all arose in the neutrally-evolving portion of the tree. However, since we want selection to come into effect for the `SLiM` phase, we will need to update the selection coefficients of these variants. Tree sequences are, for efficiency purposes, not directly modifiable. Therefore, to update any aspect of a particular tree sequence we need to instead copy its underlying tables, modify them, and then regenerate the tree sequence. Doing this to change all selection coefficients to 0.001 looks like this:

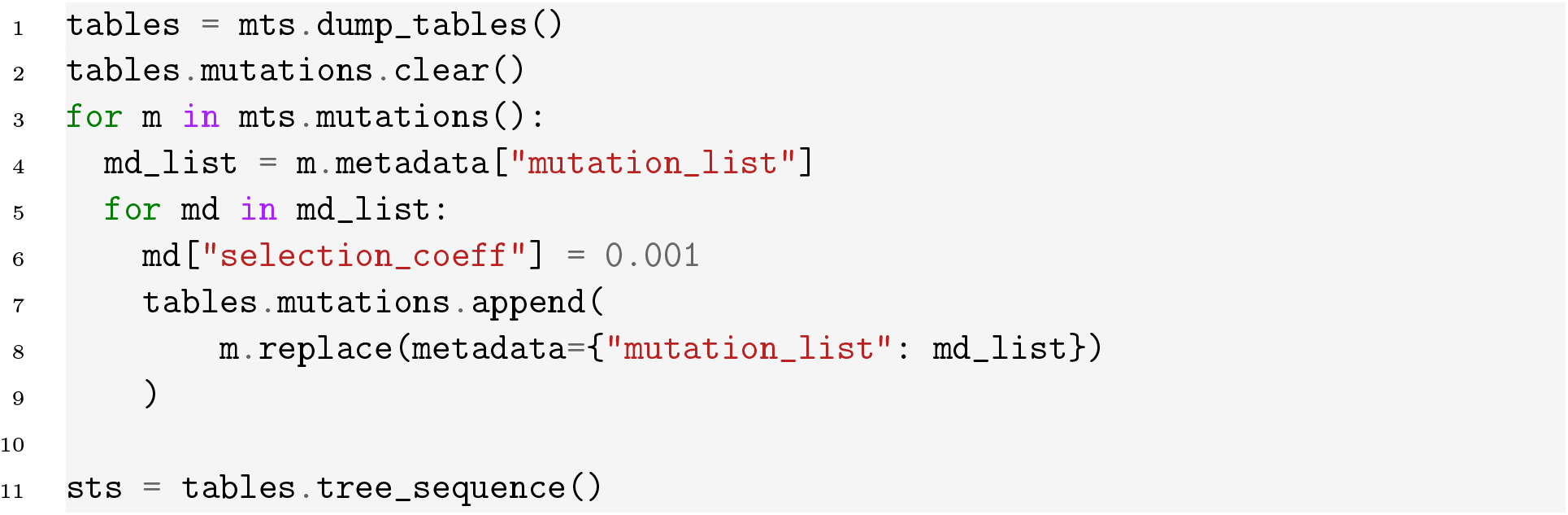

This code first copied all the tables that describe the tree sequence `mts`, cleared all the information in the mutations table, and then added each existing mutation from `mts` back with a new selection coefficient. Finally, it generated a new tree sequence from the resulting, modified, tables. This tree sequence can now be read into `SLiM` in order to proceed with the forward-in-time phase of our coalescent simulation.

Before we proceed to that step, however, we should take a detour through some details about how mutations are represented in both `SLiM` and `tskit`. This is necessary to understand the above code (what is `mutation_list`?), and it has potentially confusing implications for hybrid simulations. In `tskit`, each mutation has a “derived state”, which is the allelic state that completely replaces any previous allelic state. So, if a mutation occurs at a site where a mutation already exists, the most recent mutation will replace the older one. This differs from `SLiM`, which, by default, allows recurrent mutations at the same genomic position to ‘stack’ on top of each other. So, the `SLiMMutationModel` encodes this “stacking” information within the `tskit` mutation object. Consider the following example of a ‘double hit’ mutation that could have been produced by the code above, produced by `print(sts.mutation(295))` (it is the 622^th^ mutation in `sts`):

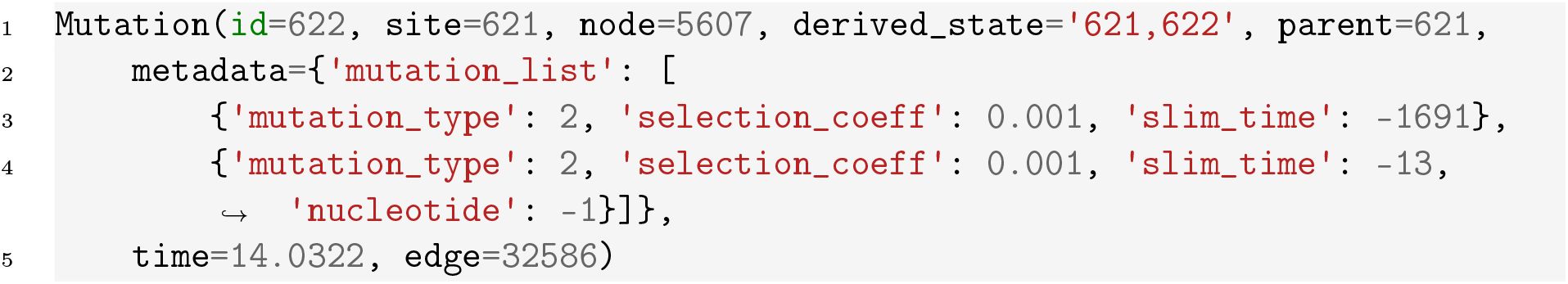

There are two parts of this record that give away the fact that this was a recurrent mutation. First, the `derived_state` is a comma-separated list of *two* integers, `621,622`. These integers are the `SLiM` IDs of the corresponding mutations. Second, the `mutation_list` element of the mutation’s metadata has two entries which represent those two `SLiM` mutations, the first (with `SLiM` ID 621) one occurring 1691 ticks ago and the second (with `SLiM` ID 622) occurring 13 ticks ago. If another mutation occurred later at this site on a genome carrying this mutation, it would be recorded as a separate mutation in the mutation tables, with derived state and mutation list having three entries each. This is why, when modifying the selection coefficients of each mutation earlier in this section, we had to have two nested loops – one iterated over mutations, while the second updated each element of the mutation’s associated `mutation_list`.

Sharp-eyed readers will note that the “time” of the mutation given by the tree sequence (14.0322 generations ago) does not exactly correspond to the `slim_time` recorded in metadata (− 13 generations in the future); in general this conversion is “rounded and sometimes off by one, depending on the model type”. For a detailed discussion of how time is converted between the tree sequence and the `SLiM` model, see the `pyslim` documentation.

### Loading into SLiM

We can finally move on to the forward-in-time component of our hybrid simulation. First, suppose we have written out the tree sequence from the previous section to file using `sts.dump(“init.trees”)`. We need to ensure that everything matches between the tree sequence and the initialization of the `SLiM` recipe we will use to load the file. Specifically, the model type must match what we specified in the tree sequence’s metadata; the genome length must be equal to the tree sequence’s `sequence_length`, and each mutation type used in the tree sequence must be declared (above, we used `m2`). In the script we will also define a placeholder mutation type `m1` for neutral mutations, to be added later. There is no need to set up a population or individuals – this is read in from the tree sequence.

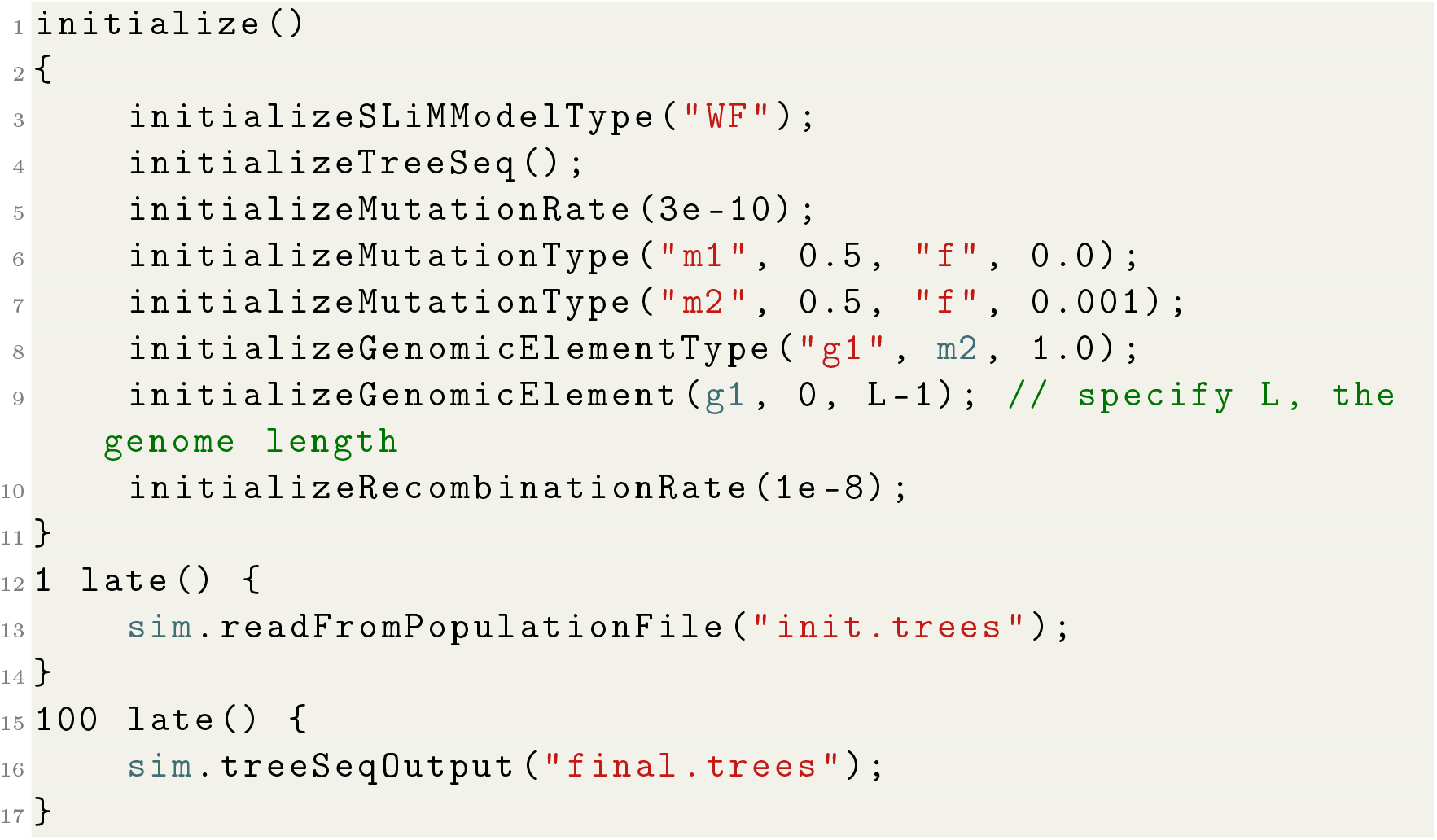

This `SLiM` code picks up the simulation where `msprime` left off, applying the same mutation rate and recombination rates for an additional 100 generations. However, in this phase, an individual’s genetic make-up (i.e., the non-neutral genetic variants they carry) will influence their fitness as the population continues to evolve.

## 4 Generating genetic data

Next, we will discuss some final steps for generating a genetic dataset from our hybrid simulation that has many of the properties of real data that is generated from sequencing or genotyping real organisms. Whether it was generated by recapitation or by first generating initial diversity with a coalescent simulation, the following steps are necessary for adding realistic levels of neutral diversity and for ultimately producing a file that can be analyzed by standard methods.

### Overlaying neutral mutations

In our two previous sections outlining the two main methods of performing a hybrid simulation, we did not bother to add neutral mutations to the tree sequence. This is because neutral mutations do not impact the shape of the genealogy, and so adding them after the fact is equivalent to tracking them as the simulation proceeds; it is more efficient because it relieves `SLiM` of the burden of tracking these irrelevant variants. In the “Generating initial diversity” section above, we stated that we wanted to simulate an overall mutation rate of 3 × 10^−8^, with 99% of the resulting mutations neutral. We already added the 1% of non-neutral mutations, so now, we need to add neutral mutations at a rate of 0.99 × 3 × 10^−8^. We will set the mutation “type” to 1 so that we can easily distinguish neutral and non-neutral mutations on this property (recall that non-neutral mutations were of type 2), matching the preamble of the `SLiM` script above. The first step is to load the tree sequence output by `SLiM`:

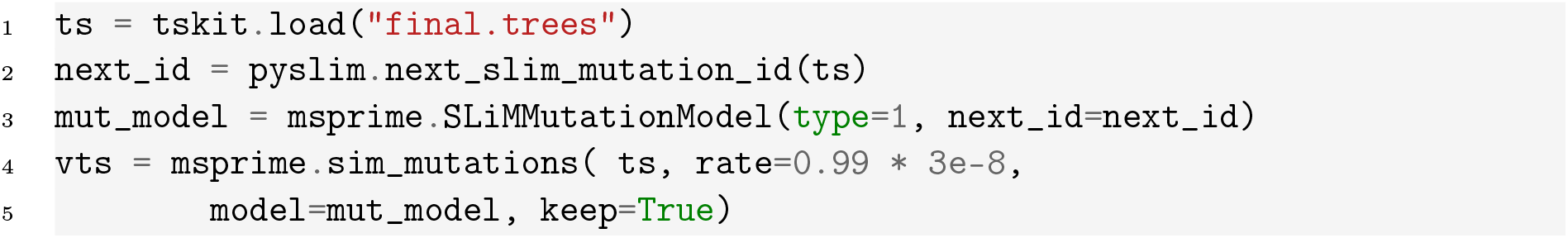

We use `pyslim`’s `next_slim_mutation_id` to identify the `SLiM` mutation ID for `msprime` to pick up from, which is important as all `SLiM` mutation IDs in the tree sequence must be unique. Additionally, we include `keep=True` so that existing mutations are not discarded.

### VCF output

The final step we demonstrate is to output the `SLiM`-coded genotype information as a Variant Call Format (VCF) file. Doing so requires remapping some of the information to a form that resembles real genetic data. For example, the `SLiM` mutation model encodes the ancestral state of every site as the empty string and the derived state as a comma-separated list of integers (also stored as a string). These states are meaningful for `SLiM` but do not correspond to the nucleotides expected in a VCF. To address this, `pyslim` provides functions to convert `SLiM` mutations to a more conventional format: first, `generate_nucleotides` adds nucleotides to `SLiM` mutations not containing nucleotides, and second, `convert_alleles` replaces the comma-separated lists of integers in the ancestral and derived states of the tree sequence with the corresponding nucleotides. Following these steps, `tskit`’s method `write_vcf` can be used to output the VCF (and the method itself provides more options):

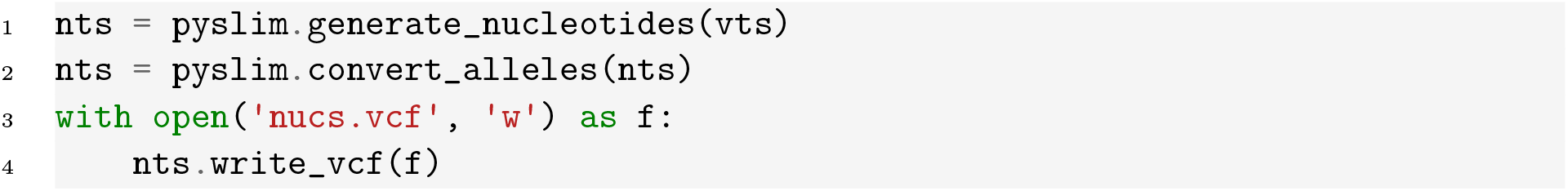

This simply assigns each position a randomly chosen nucleotide, and then assigns each mutation a nucleotide chosen randomly out of the three that are different from the previous state (i.e., from the Jukes-Cantor model). More complex models could be implemented in python.

The key step in both this and the next section will be to combine the results of different simulation runs. For instance, the tree sequences produced by the simulations of branches *A* and *B* will share history up until their branching point; which we need to take into account when we put these two tree sequences back together. Since the two share some history but we want to combine the bits of history they don’t share, the operation is called “union”. Although this operation abstractly sounds straightforward – “we just need to take the bits of the tree sequence that happens on the branch to *B* and add it to the tree sequence from *A*” – in practice many things can go wrong. To ensure that we can put them together smoothly, we need to make sure that the parts of shared history recorded in each simulation agree. This is discussed in more detail below.

## 5 Parallelizing forward-in-time simulations of multiple species

We have thus far demonstrated how to combine backwards- and forwards-in-time simulations in two different ways. Now, we will discuss how to distribute the computational load of a highly complex forward-in-time simulation across multiple processes operating in parallel, by simulating different branches of a phylogeny independently and stitching the resulting tree sequences together. This approach can drastically reduce overall clock time and the memory requirements of a large phylogenetic simulation. The main challenge with this approach then becomes determining how to stitch together multiple ARGs that represent the same time span.

The idea works because if there is no migration between species, any two branches stemming from the same node in a given tree do not affect each other, and thus can be simulated in parallel. Figure 3 shows the simple example we work with.

**Figure 3.**
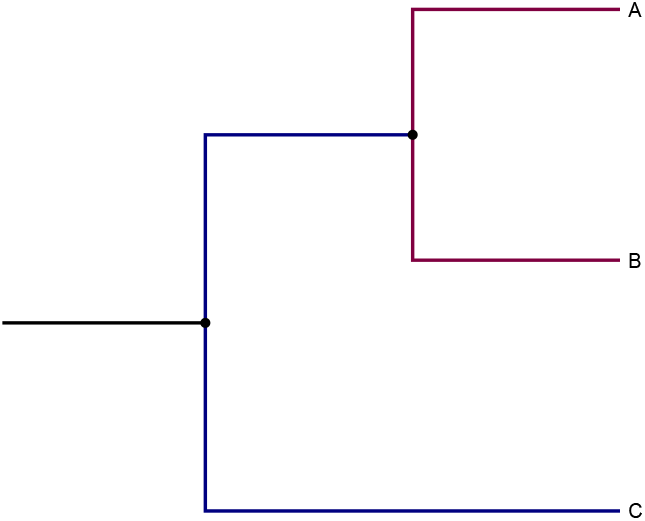
Example phylogeny. Branches with the same color can be simulated in parallel if there is no migration between them.

### 5.1 Parallel simulation of branches

We will use the following `SLiM` script to simulate the history of each species, represented by its own branch, in our phylogeny. Here, each species can have a different (but fixed) population size (`popsize`) and total duration (`num_gen`). Additionally, we will simulate deleterious mutations accumulating across the entire genome at a fixed rate.

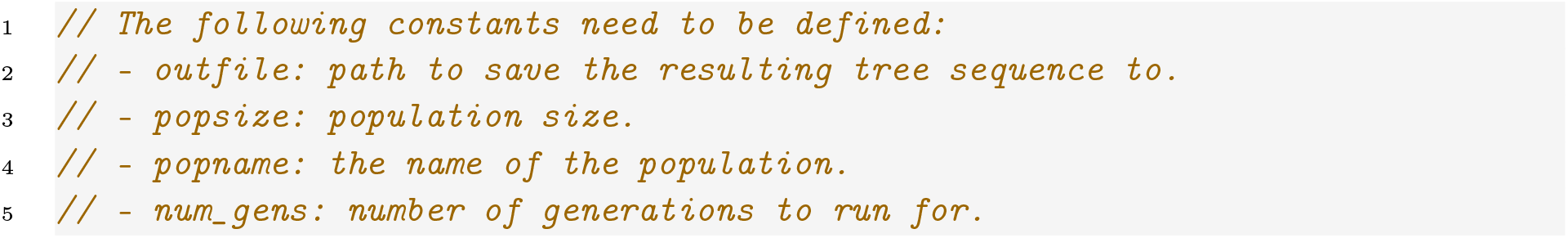

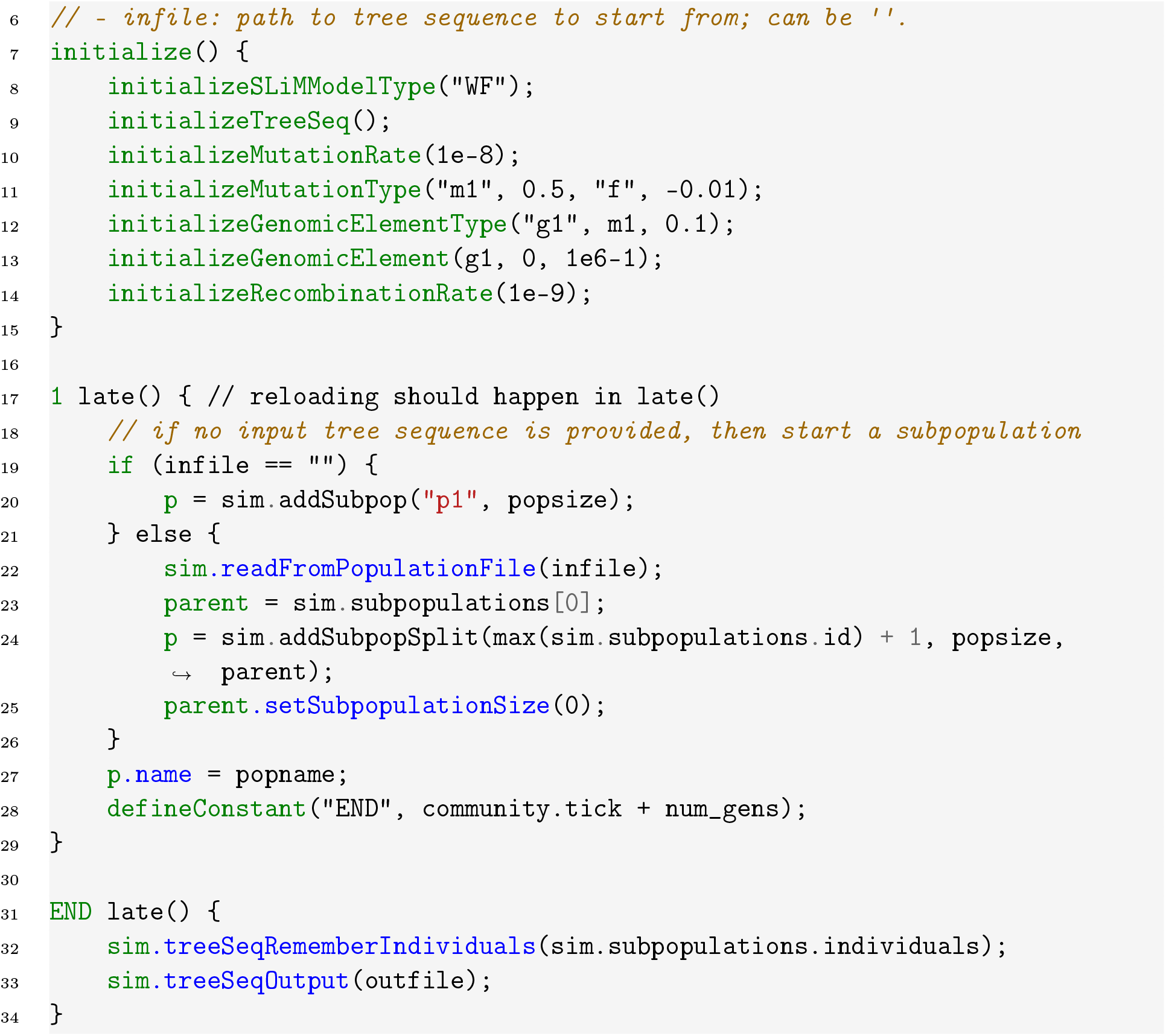

Notice that, in addition to `popsize`, `popname` and `num_gen`, the variables `infile` and `outfile` are left undefined in this `SLiM` script. This allows the relationship between branches, and the resulting tree sequences, to be explicitly defined. As a result, this single `SLiM` model can represent every species simply by passing the appropriate values of `infile` and `outfile`. When initializing a `SLiM` simulation using an existing tree sequence, the starting tick is updated based on the time specified in the tree sequence’s metadata, as seen above. Therefore, we define the ending tick of the simulation as the start time plus the `num_gens` argument.

At the end of the simulation, we call `sim.treeSeqRememberIndividuals` right before saving the resulting tree sequence. This is necessary to ensure that the individuals in the final generation are never dropped from any of the future runs that are started from this simulation’s output, as they will be needed later to merge the tree sequences generated from parallel runs back together.

To actually run our parallelized simulation, we could use workflow management software, such as Make or Snakemake, or a custom script – see the `pyslim` documentation for details on different implementations. Here, we will use the simple Make script below to pass the appropriate parameters to our generic `simulate_branch.slim` script.

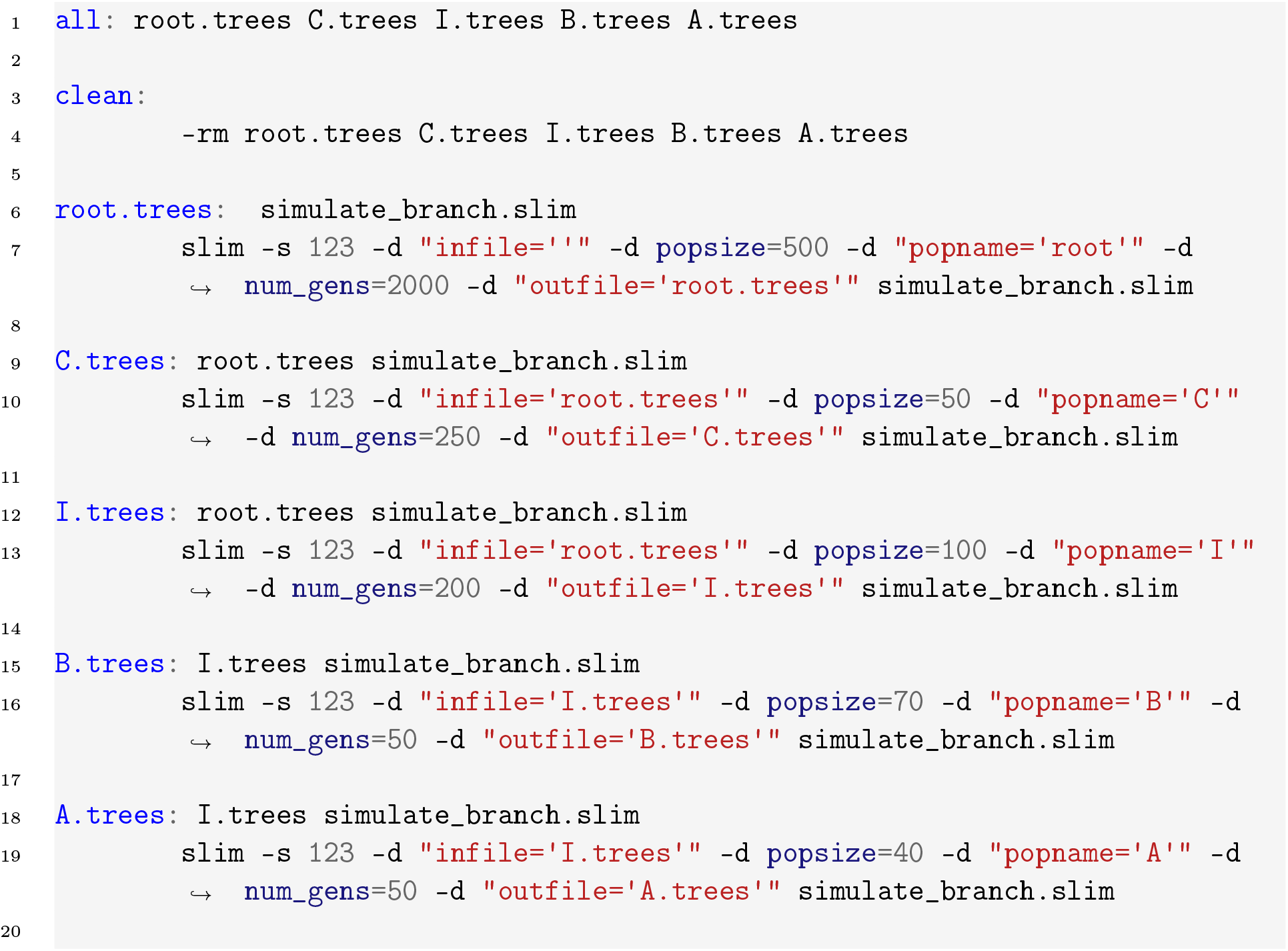

This script essentially recodes our original phylogeny through a list of ‘rules’ that define each parent-child relationship. For example, the files ‘A.trees’ and ‘B.trees’ both depend on the internal node ‘I.trees’, which in turn depends on the result of simulating the ultimate ancestral branch ‘root.trees’. Each rule in the Makefile also specifies how to generate the output given the required input through a single line of `SLiM` code.

We can then run these simulations in parallel by executing the following bash code, in which the -j argument specifies the maximum number of simulations to run simultaneously:

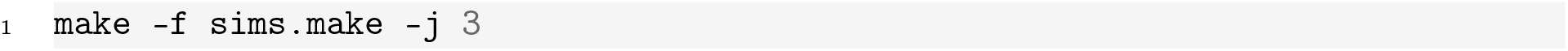

### 5.2 Putting it back together: unioning the tree sequences

We now need to stitch the resulting tree sequences back together. Remember, each of the tree sequences representing species ‘A’, ‘B’, and ‘C’ are distinct, but overlap significantly as a result of their shared evolutionary history. To join them together, we need to specify which bits are shared and which are not. This can be accomplished for pairs of tree sequences using the `union` operation from `tskit`, which simply copies the unique parts of one of the tree sequence onto the other. Figure 4 illustrates how this works.

**Figure 4.**
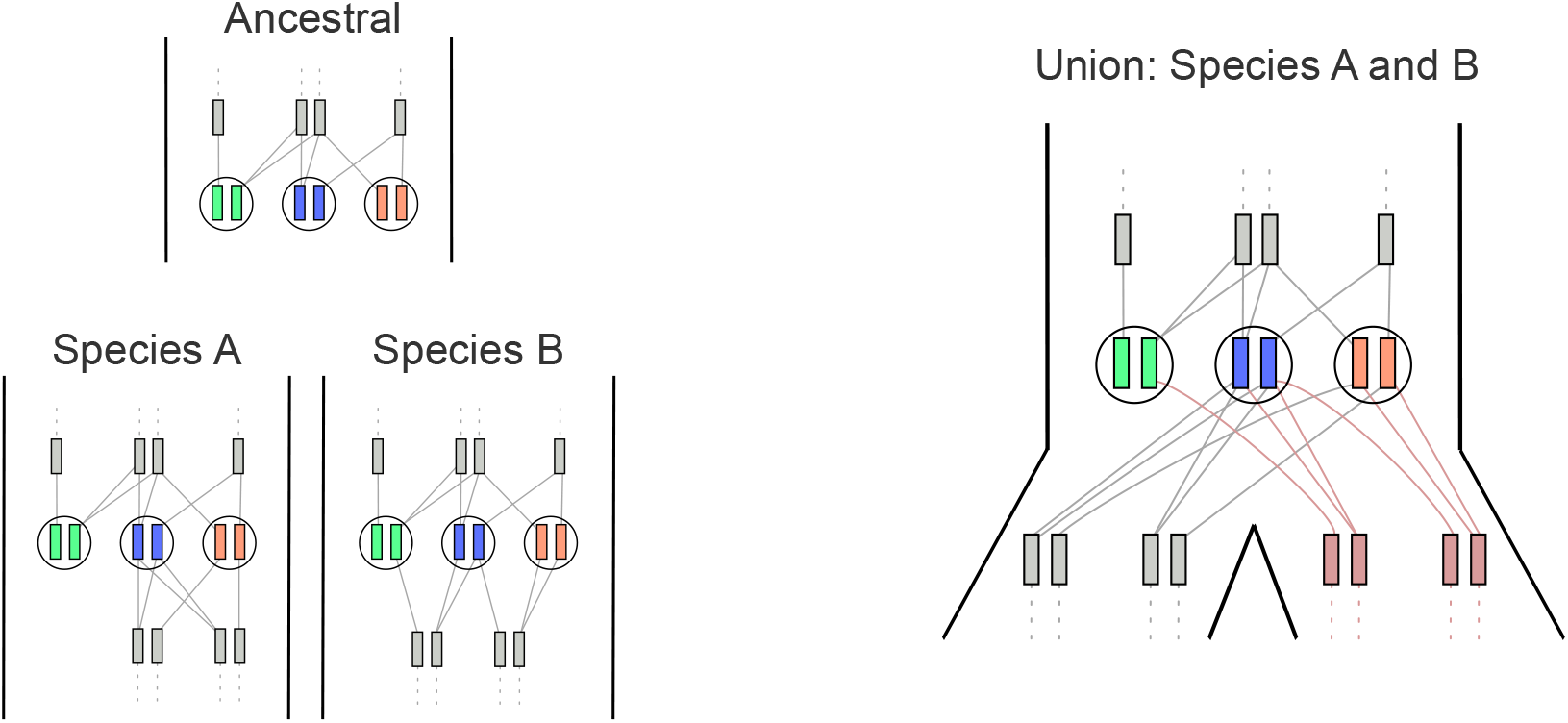
Species A and B share the same ancestral history, but after the split they are simulated independently. Note how the history above the split (delimited by the individuals which are remembered and persistent across tree sequences, highlighted in green, blue and orange) is identical, thanks to the call to treeSeqRememberIndividuals at the end of each branch. tskit’s union operation will merge these tree sequences given a mapping that communicates which nodes in the two tree sequences are the same. In this example, union would add to A any new history found in B; these new nodes and relationships are shown in light red in the result.

The key to correctly combining a given pair of tree sequences is to identify which parts of each are shared between the two. There are potentially many ways to do this; `union` works by identifying nodes, and so requires a ‘map’ that matches shared nodes across the two tree sequences (and unmatched nodes are new). Using this map, `union` (a) checks that any properties of shared nodes and relationships between them are identical, and (b) appends any new nodes and relationships found in the second tree sequence to the first one. Luckily, `SLiM` stores a value called `slim_id` in the metadata of each node that it produces, which we can use to identify the shared nodes. However, a matching `slim_id` is not sufficient to identify a positive match, because although `slim_id` values are unique within a given `SLiM` simulation, the same IDs will be used in different (parallel) simulations. When `SLiM` loads a tree sequence in order to continue a simulation, it will start assigning `slim_id` values to new nodes using the last `slim_id` value present in the ancestral tree sequence as a starting point. So, when more than one simulation is initiated with the same ancestral tree sequence and the resulting tree sequences are merged back together, it is likely for the same `slim_id` to be used to refer to different nodes. However, we can use node *time* to resolve the issue: referring to Figure 4, if two nodes that were born at or before the split time between ‘A’ and ‘B’ have the same `slim_id`, then they must represent the same ancestral genome (and so must have exactly the same time); while if they were born after the split time, they represent different ancestral genomes. The following function constructs a “node map” that identifies the nodes shared by two tree sequences, `other` and `ts`, by identifying pairs of nodes that arose before `split_time` and have the same `slim_id`. The output is an array with one value for each node in `other`, whose `j`th entry is the ID of the matching node in `ts` if the node is shared, and is `-1` otherwise. This code could be made easier to read by iterating over nodes, but the following numpy-based version is much faster:

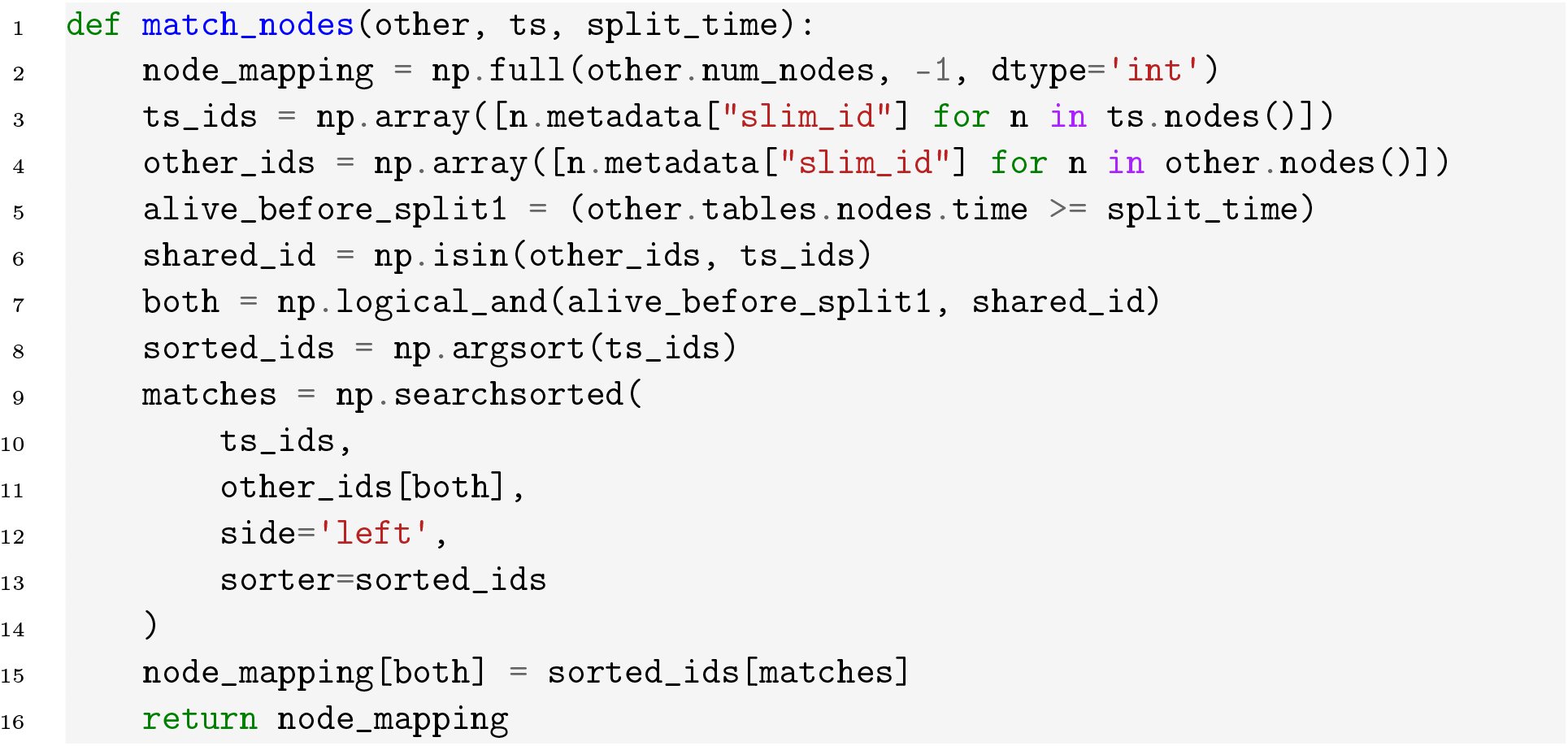

This node map can then be used with `ts.union` to combine our tree sequences pairwise. A general-purpose recursive strategy to do this is provided in the `pyslim` documentation, but the problem is easy in this simple example:

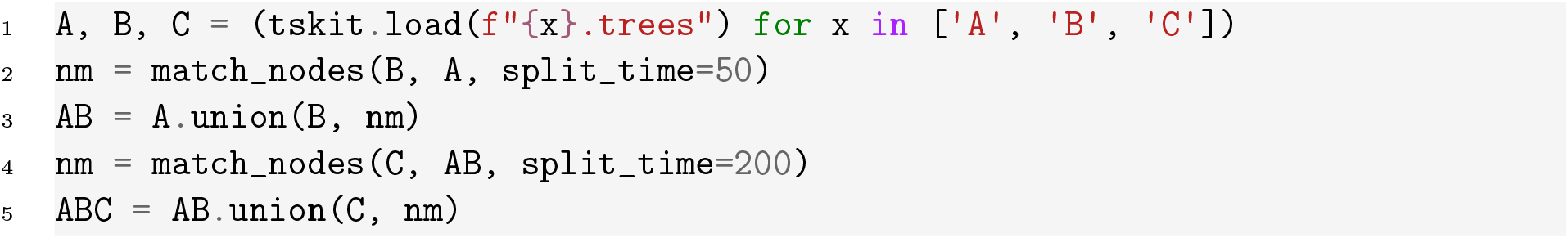

The result of executing this code is a single tree sequence, `ABC`, that fully represents the genealogy of the sampled individuals from all three extant species in our phylogeny (Figure 3). Because each branch of the tree was modeled as a distinct population and `SLiM`, it is straightforward to identify which nodes and individuals belong to which branch on the phylogeny throughout time. After potentially recapitating and adding neutral mutations, we could analyze this tree sequence like any other to answer a variety of questions related to the evolutionary history and genetic ancestry of these species.

When adapting this code for your own use, it is common to run into a situation where you get an error saying that “shared portions of the tree sequences are not equal”. Although it is possible to, it is important to *not* disable this check: almost certainly, this error is alerting you to a problem, and proceeding with the union would produce incorrect results.

The major advantage of running simulations in parallel is that it expands the scope for complexity and size of feasible models. We can simulate many more species, or much larger populations within those species, if we can split the computational burden in this way. For example, Rodrigues et al. [2024] used this parallelized approach to conduct whole-chromosome simulations of the entire history of great apes spanning over 10 million years. In this case, some individual species simulations used over 100Gb of memory, and so without parallelization, simulating great apes species history in a single process would have been effectively impossible given the current limitations on computational resources.

## 6 Meta-population dynamics with simulation networks

Finally, we consider a more complex scenario that requires more advanced manipulations of the tree sequence: simulation of the within-host dynamics of a pathogen, where individual hosts are simulated in parallel using the same basic `SLiM` script. This is therefore similar to the previous example, but more complex for two reasons: 1) we don’t want to retain the entire population at branch points, for efficiency reasons; and 2) we will allow for reticulations in the “phylogeny”, which occur when a host individual is infected from more than one source.

We can model a pathogen infecting a host species as a large meta-population, where each infected host constitutes a distinct pathogen deme. Similar to our previous section, we can represent the relationship between individual hosts with a phylogeny-like branching structure, as in Figure 5. In this figure, nodes represent pathogen subpopulations within individual hosts and transmission events from one host to another are represented by edges.

**Figure 5.**
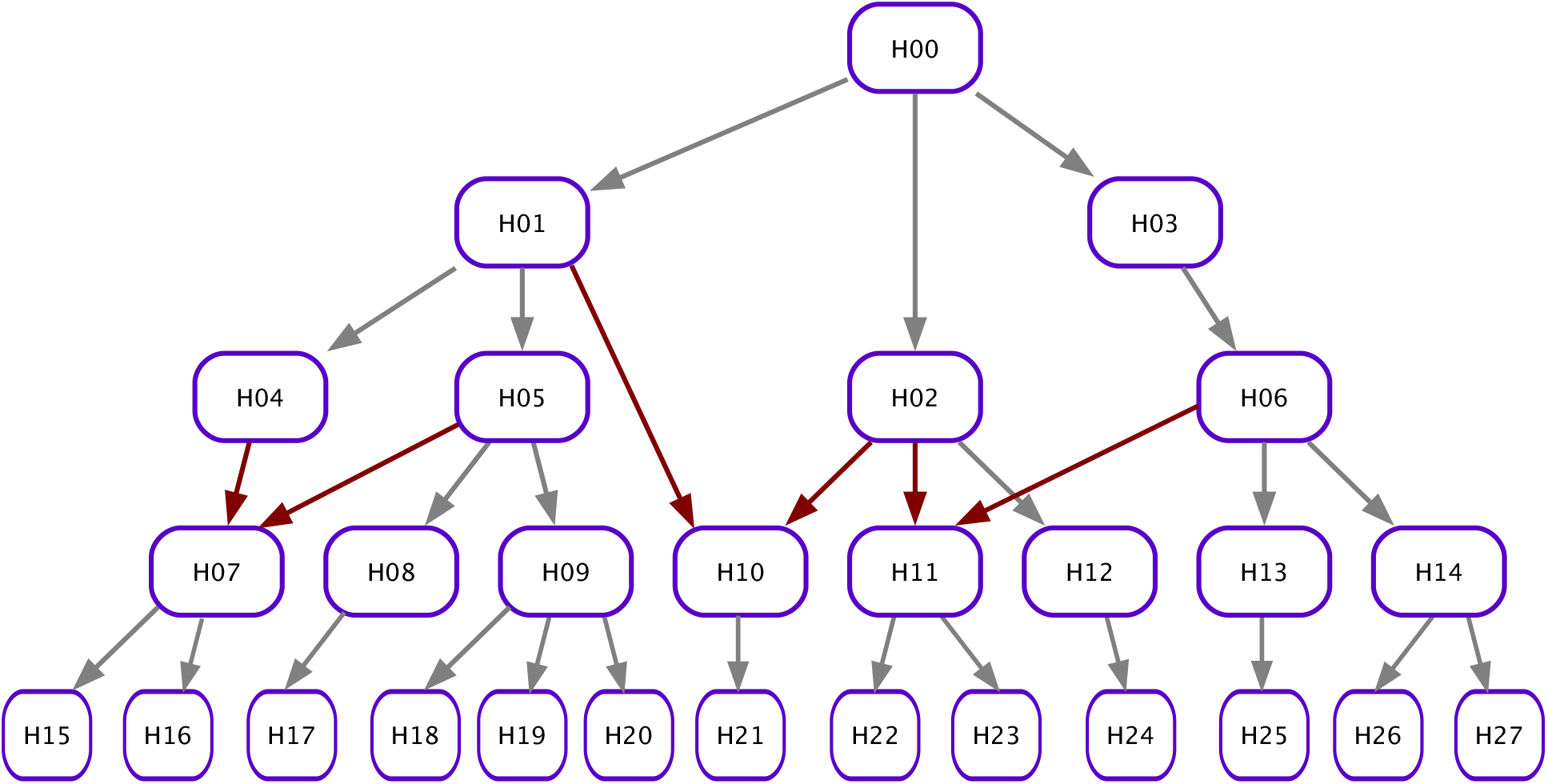
Nodes in this figure represent individual host infections, while the arrows represent the direction in which pathogens are transmitted from one to the next. Here, red arrows represent cases of reticulation in the transmission graph, where a host infection is founded by pathogens from two distinct source infections.

Just as in the prior vignette, the backbone of this model is a flexible `SLiM` script that can accept a specific tree sequence as a starting point in addition to other parameter values. This script simulates the logistic growth of pathogen individuals within a host, as well as a bottleneck of a few pathogens to be transmitted to each subsequent host. These dynamics will play out repeatedly as we model the pathogen spreading through the host population.

Three major differences to the previous model are that branching (i.e., transmission events) can occur at various times during a given host infection, that one simulation can begin with the results of more than one previous simulation, and that we use a non-Wright-Fisher model with overlapping generations. Based on the previous approach, a first try at this might be to save the tree sequence at the times of each new infection. However, this fails: previously, we could assume when combining two simulations (see Figure 4) that all shared individuals had identical lives, but with overlapping generations more care must be taken. If two simulations begin with the same tree sequence for an initial state, then the “same” individuals could live parallel and contradictory lives along the two branches of the simulation. Furthermore, we would like to save only those few individuals who start each new infection, rather than the entire population. So, we will store only the information needed for piecing the full genealogy back together, allowing `SLiM` to simplify out the rest to maintain efficiency. Each host simulation produces a single output tree sequence that contains all the samples required to represent all downstream transmission events. Then, we make use of advanced operations to use this single tree sequence to start several different downstream simulations.

The basic workflow for a within-host `SLiM` simulation is as follows. The infection starts by loading an input tree sequence, if specified, as well as additional information that specifies which hosts the current infection will be transmitted to (`OUTPUT_HOST_IDS`), when these transmission events will happen, relative to the start of the simulation, (`TRANSMISSION_DAYS`), and how many pathogen individuals will be transmitted (`NUM_SAMPLES`). The simulation proceeds with the pathogen reproducing according to some rules; in our case, we parametrize this process by setting an overall growth rate (`POP_GROWTH_RATE`) and a carrying capacity for pathogens within the host (`K`).

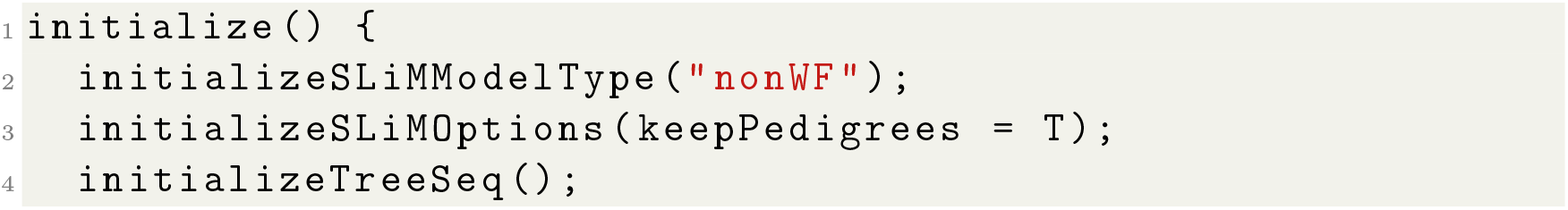
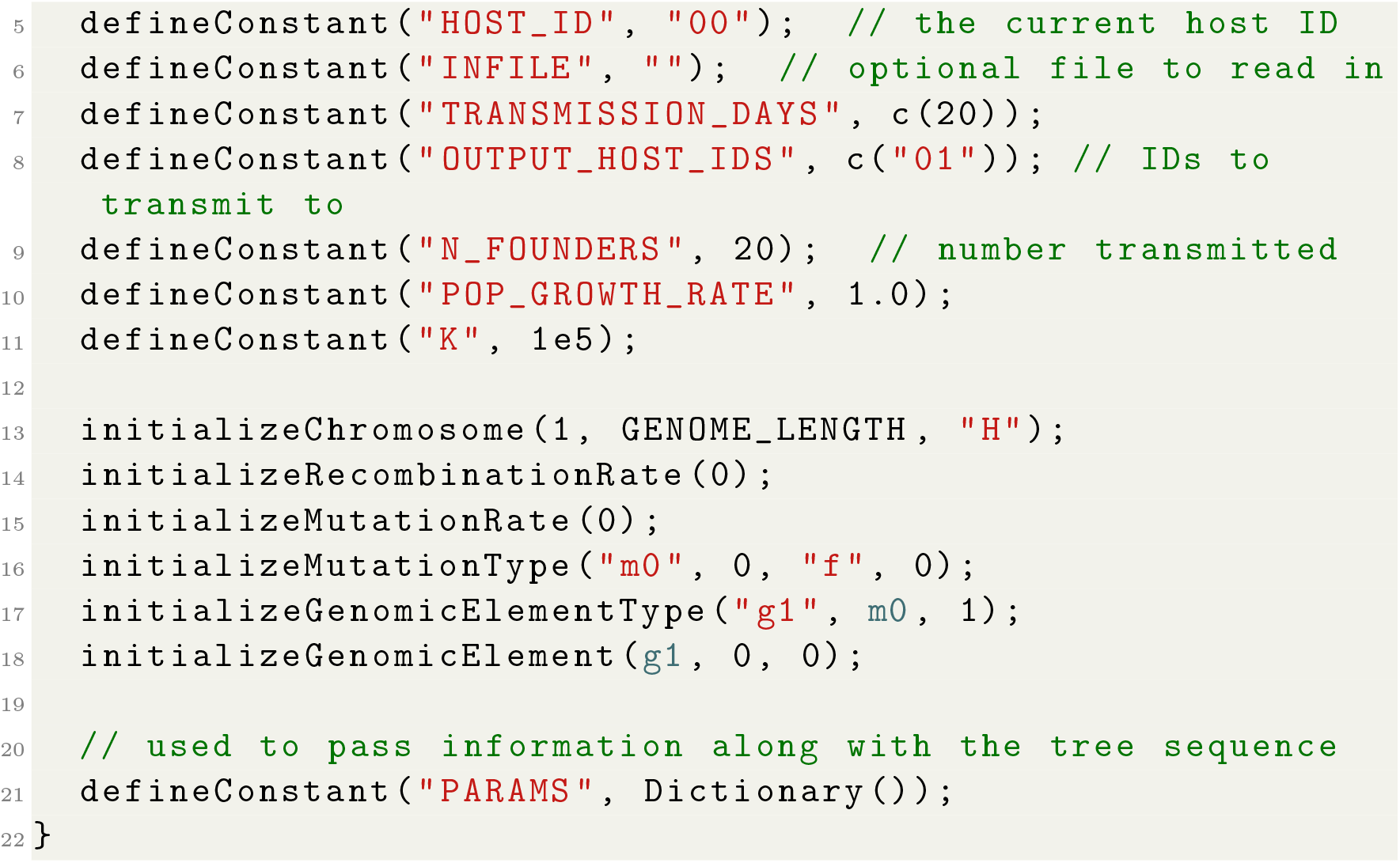

If there is no input file given, we begin by creating a new subpopulation in `SLiM` whose ID is equal to the current host ID. If there is an input file, this subpopulation is created upon loading that file. This step also creates dictionaries for founder pathogen IDs and infection start times (i.e., *founding times*) that will be stored later in metadata, for access within `SLiM` as well as `tskit`. If there is a tree sequence file given, we reload the simulation state from the file, as well as additional *metadata* from the tree sequence file. In the metadata, we’ve stored the IDs of the individuals who are to be the founders, so we move these to a newly created subpopulation. Finally, this code calculates the *absolute transmission times* (the day of the overall simulation) by adding the start time of the current simulation to transmission days supplied.

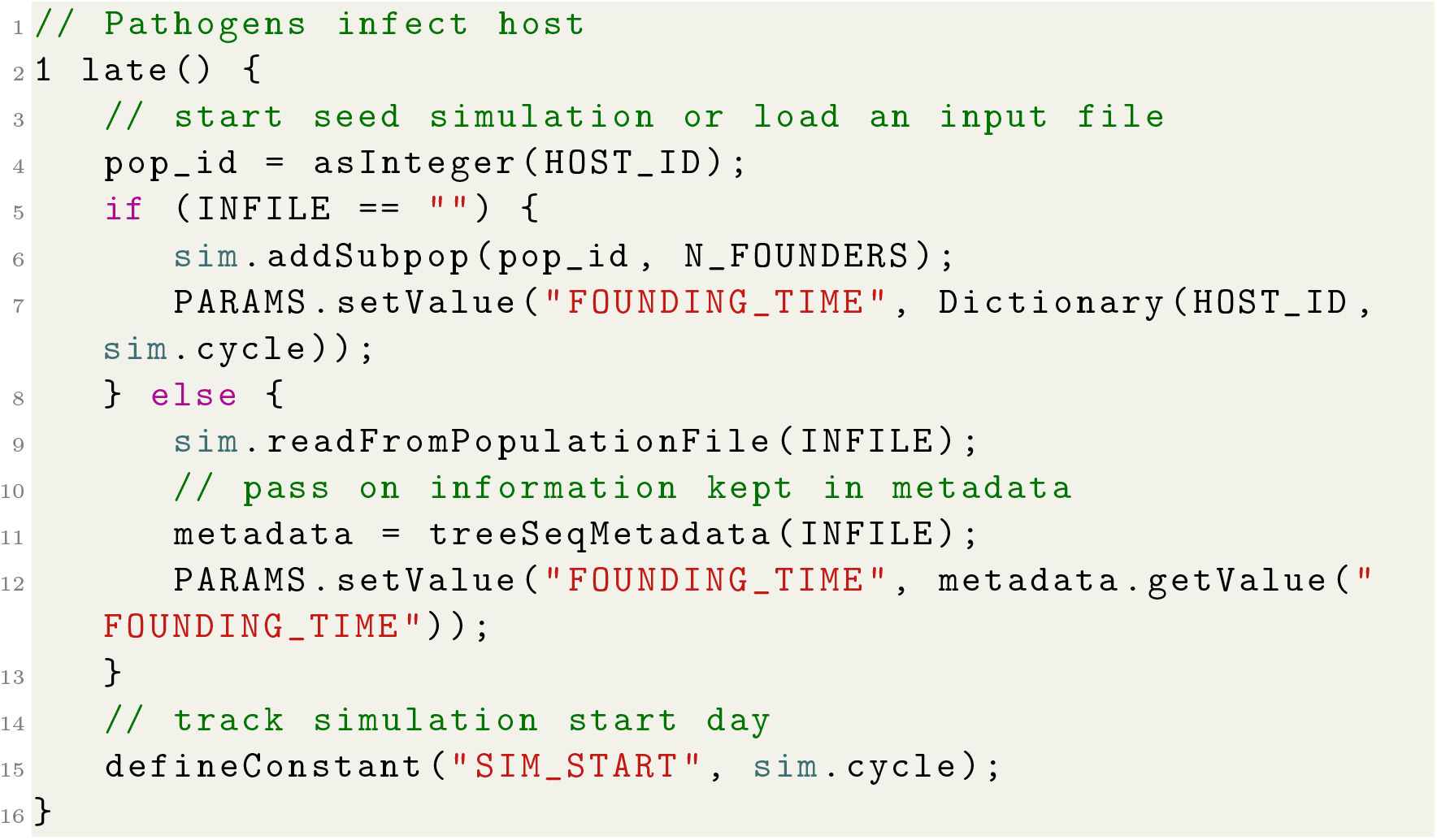

We then allow this population to evolve. Each individual produces a random number of clonal offspring. Then, every individual’s probability of survival (their `fitnessScaling`) is set depending on the overall population size; when the population exceeds the carrying capacity, probability of survival is reduced to bring their numbers back down.

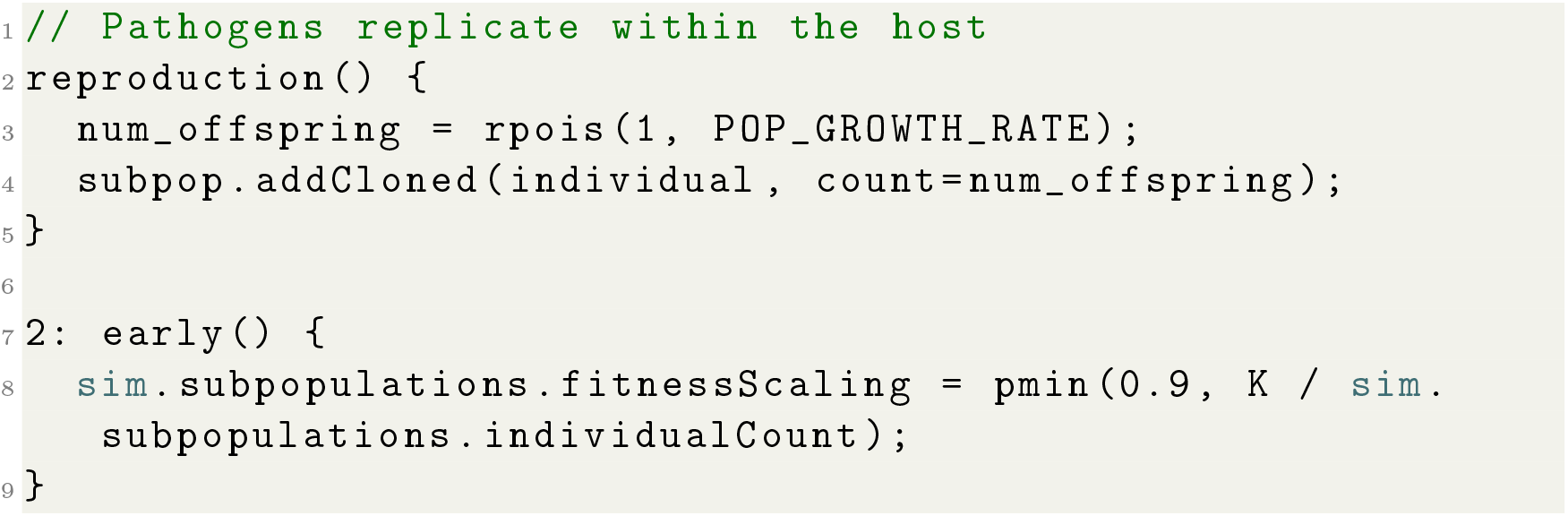

When the simulation reaches one of the designated *transmission days*, an additional callback is executed that transmits some individuals to a subsequent host. To do this, the code first identifies which host IDs should be transmitted to, and then randomly chooses which individuals will be the “founders” for these new infections. Each founder is then moved to a new subpopulation whose ID matches the ID of the new host ID, remembered, and killed. (Recall that “remembering” an individual means that they remain in the tree sequence forever.) As we will see, these individuals are only dead temporarily; they will be resurrected in order to continue the simulation as the founders of the next host infection. A final piece of bookkeeping is to store information about this event in metadata: the `FOUNDING_TIME` is a dictionary that simply records when this transmission event for each host happens. This information (along with everything else in `PARAMS`) is attached to the output tree sequence as top-level metadata. This property of tree sequences, which is accessible in both `SLiM` and python, is extremely useful because it can store arbitrary bits of information that we might want later. In our case, we will use `FOUNDING_TIME` to determine how to properly resurrect these founding individuals.

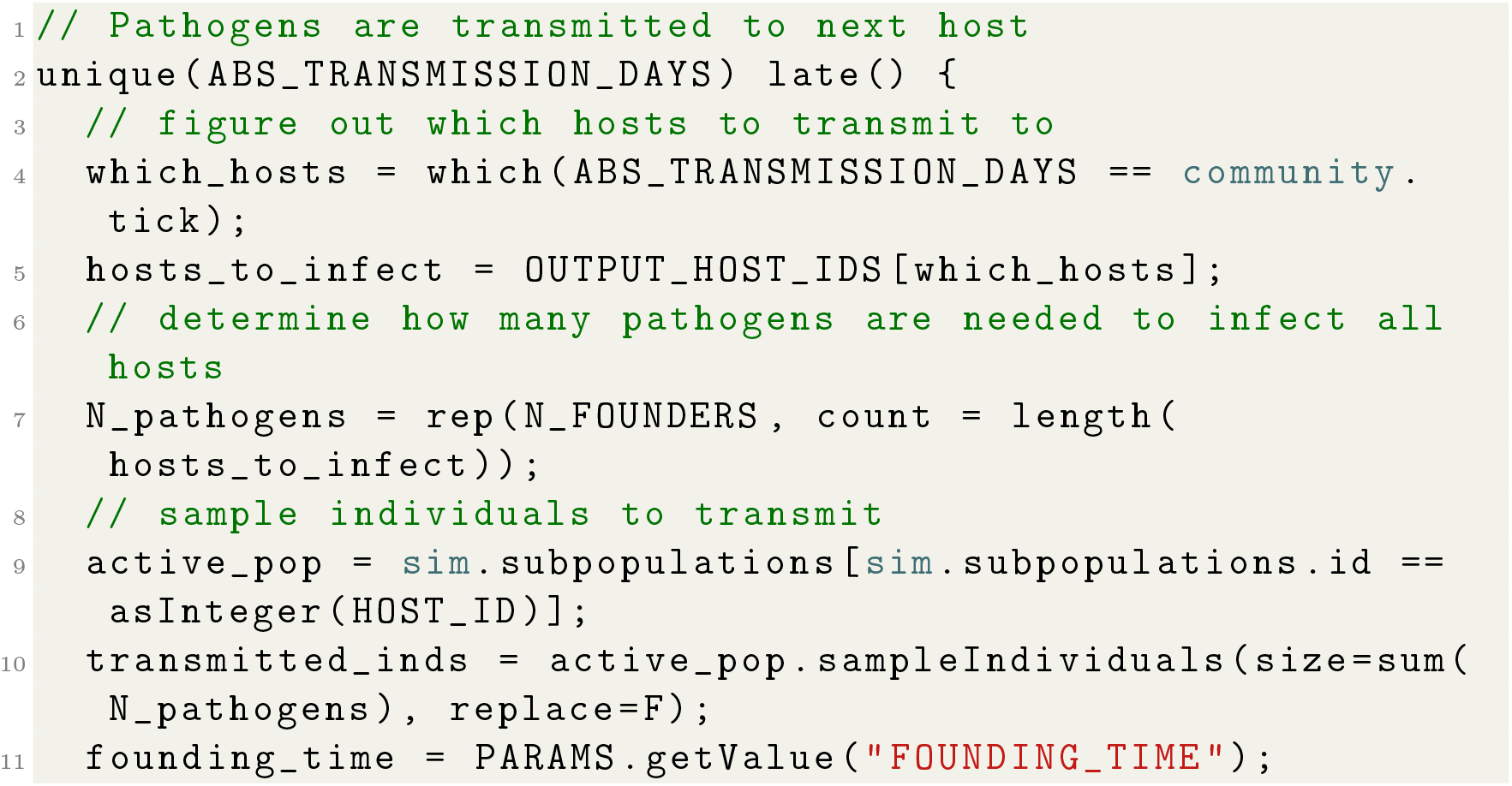
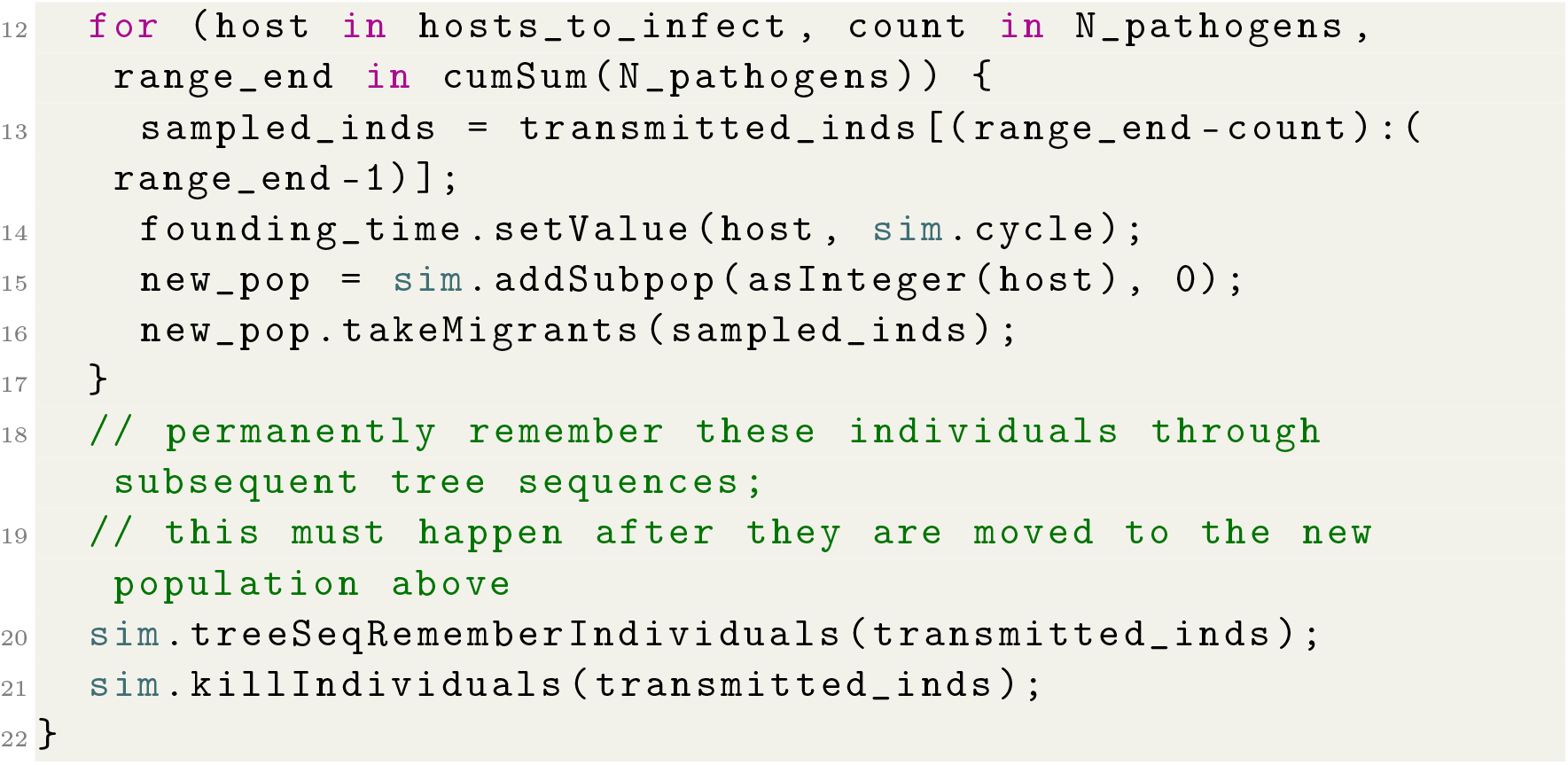

This simulation runs as long as necessary to complete all the specified transmission events, and then ends by writing out a single output tree sequence. In this tree sequence, all individuals that were transmitted at any point in the simulation will be *remembered* as samples, but will be dead. Additionally, we kill all remaining individuals in the host population just before writing the output. If we did not do this, thousands of individuals alive at the final tick of the simulation would also be marked as samples and their full genealogical history would be stored in the tree sequence. Because we only care about the relatively few individuals that will go on to found new infections, we can save significant time and memory by getting rid of everyone else, too.

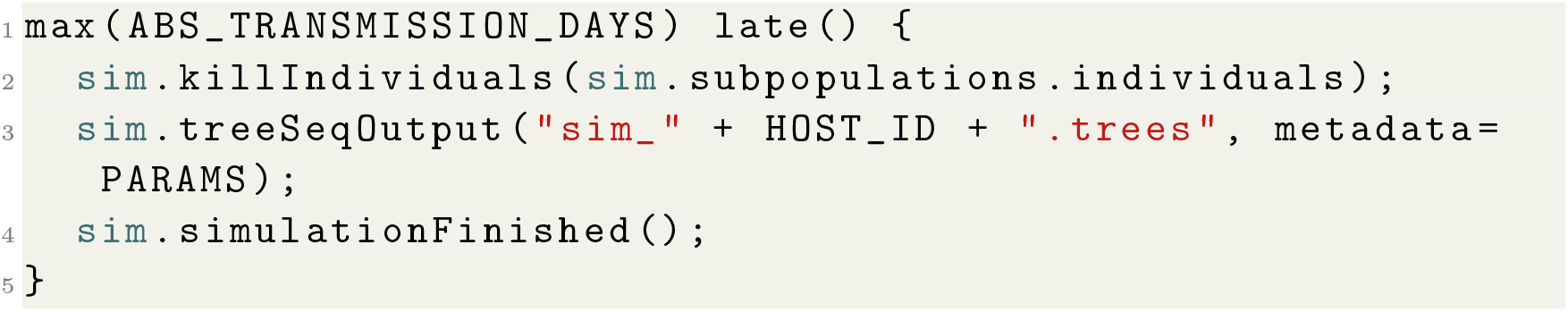

Now we have a tree sequence with no living individuals at all, which will not be of much use if we were to load it back into `SLiM` directly. We therefore need to do some processing with `tskit` before we can initiate the next set of infections, which is accomplished with a python script. This script is passed a tree sequence and a host ID, and then retrieves the starting time of that respective infection from top-level metadata (from `FOUNDING_TIME`). The script then uses the `set_slim_state` function in pyslim to rewind the tree sequence’s time, and also to resurrect any individuals that were still alive at that time. This function does two things: it resets the time as recorded by `SLiM` in the tree sequence, and labels a given set of individuals as being alive. This results in a curious thing: the tree sequence can contain information about the “future”. Of course, `SLiM` only pays attention to those individuals that are marked as being alive, so this is harmless. The argument to `set_slim_state` is in `tskit` time ago, while the value we stored in metadata was a `SLiM` tick, so to convert we subtract the stored tick from the current tick, which we obtain from top-level metadata. This step is done for every host that the current infection transmits to, producing a distinct version of the output tree sequence for each one.

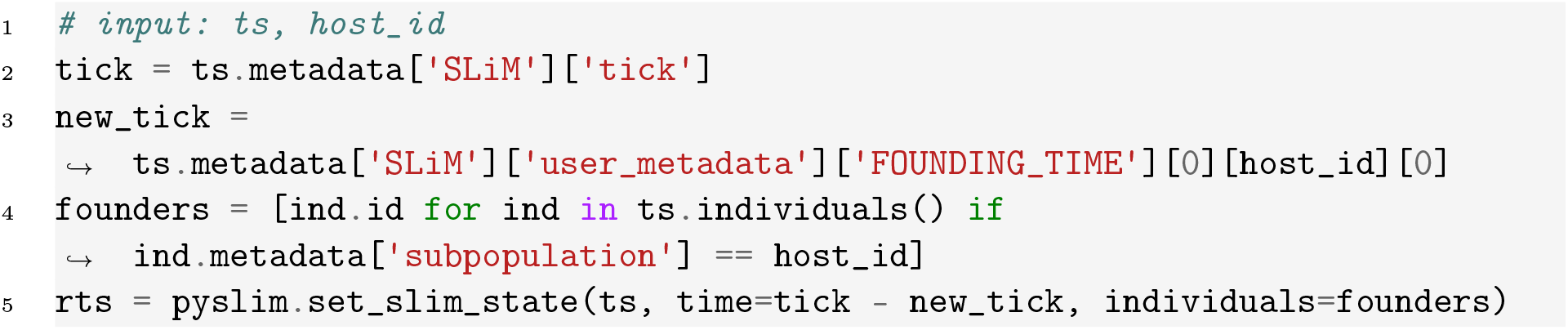

Then, `rts` is written out, to start a new simulation. The code above is somewhat simplified for clarity: in practice we also remove individuals with duplicate pedigree IDs, which can happen when the input tree sequence was produced by merging the output from two distinct simulations.

For hosts who are infected by founders from more than one other infection, before doing the previous step we need to merge the tree sequences that are output by the upstream simulations. For instance, if host 4 is infected by pathogens from both hosts 1 and 2, then the tree sequences output from both hosts 1 and 2 have founders we need to start the simulation for host 4. So, we need to merge these (using `union`). We will also want to merge the multiple tree sequences produced by all terminal simulations into one as well. In our previous example, we used the split time between two branches to disambiguate distinct nodes that might happen to share a `slim_id`. That won’t work here because some of the shared nodes can be more recent than the split time, thanks to the resetting done by `set_slim_state`. As previously noted, we use the current host ID as the subpopulation ID within `SLiM`. Since host IDs are unique, we can use a combination of `slim_id` and `population` to definitively match them across pairs of tree sequences, and to generate the node map required by `tskit.union`. Tree sequence merging can then be achieved in the same way as outlined in our previous vignette.

However, there are a few additional steps. These are required basically because we need to help `union` identify what is “the same” between tree sequences and what is not. So, the python script that merges two tree sequences: (1) Shifts the node and mutation times in each input so that time matches, by adding to the “time ago” of each the difference between their `SLiM` tick and the largest `SLiM` tick. (2) Merges the population tables so that any populations described in the second are also described in the first. (3) Updates individual metadata and flags for any individuals contained in both, identified by unique combinations of birth `population`, `pedigree_id`, so that these agree between the two tree sequences for shared individuals. This is necessary for rare cases in which, for instance, a founder of one host infection is then passed on to found yet another infection in another host, in which case the age and subpopulation recorded in metadata in the two tree sequences will not match (and we want to use the later one). (4) Constructs a node mapping from the second to the first, by identifying those nodes sharing unique (birth `population`, `slim_id`) pairs. (5) Unions the two with this mapping. (6) Copies missing keys from `FOUNDING_TIME` in the second’s metadata to the union.

Similar to the previous example, we can run our simulation in parallel using a job scheduler like Make. The framework we outline here makes it possible to model the evolution of very large and complex pathogen meta-populations in order to understand how both among-host and within-host factors impact these processes. For example, we can use our model to develop expectations for how pathogens evolve under variable selection pressures. Imagine a life history trade-off between replication and transmissibility, so that one allele at a particular locus allows the individual carrying it to reproduce at a faster rate while the alternate allele increases an individual’s likelihood of being transmitted to a new host. Similar trade-offs have been observed in influenza, HIV, and COVID-19 [Liang, 2023, Ariën et al., 2005, Zhu et al., 2022], but would be challenging to model realistically without parallelization.

## Discussion

Moving from simulation of genotype data to full ARGs presents many opportunities for evolutionary genomic inference. ARGs can store a great deal more information about a population’s evolutionary history, while making simulation more efficient [Kelleher et al., 2018]. Furthermore, as we discuss here, information can also be passed between different simulations via the ARG, which enables hybrid simulations that combine both forward- and backward-in-time components as well as parallel simulations. In this chapter, we have sought to introduce readers to these methods’ potential to expand the scope of evolutionary simulation. Additionally, we have provided practical vignettes that use `pyslim`, a software package that provides functions to make model interoperability between `SLiM` and `tskit` more seamless.

In this paper, we have opted to cover opportunities and strategies more than technical detail, and so for some technical aspects, advanced users will need to consult the documentation. One important aspect is *time units*: we alluded to the fact that time units in `SLiM` and `tskit` differ in direction, and that exact conversion between them can be difficult: it depends on the stage in the `SLiM` model. More importantly, for non-Wright-Fisher models in `SLiM`, a single tick may not necessarily equal one generation, so recombination or mutation rates may need to be adjusted. The topic is covered in more detail in the `pyslim` documentation.

An overarching theme here, and for evolutionary simulations more generally, is the trade-off between realism and efficiency. Forward-in-time simulations can be fashioned to more naturally match real population dynamics, but this generally comes at the cost of speed. On the other hand, coalescent simulations make many assumptions, many of which are unrealistic, in exchange for faster runtimes. The same trade-off occurs when considering rescaling [Cury et al., 2022, Dabi and Schrider, 2025]. In this paper, we describe how users can begin to achieve the best of both worlds, outlining several examples that leverage hybrid approaches and parallelization to model scenarios that would otherwise be computationally intractable.

While approximations are always necessary in modeling, advances in simulation and ARG software have gone a long way to increase realism. For example, we discuss how we can use a coalescent phase to reduce the burden of simulating a burn-in period. As a result, ancient events occur under a much less realistic model, but this may be acceptable in certain situations. As with all cases where the validity of a particular approximation may be questionable, a good strategy is to adjust its parameters (e.g., the duration of the forward-in-time phase prior to recapitation) to see if important conclusions are affected.

